# Comprehensive analysis of behavioral dynamics in the protochordate *Ciona intestinalis*

**DOI:** 10.1101/2021.10.29.466420

**Authors:** Athira Athira, Daniel Dondorp, Jerneja Rudolf, Olivia Peytral, Marios Chatzigeorgiou

**Affiliations:** Sars International Centre for Marine Molecular Biology, University of Bergen, 5006, Norway

## Abstract

Vertebrate nervous systems can generate a remarkable diversity of behaviors. However, our understanding of how behaviors may have evolved in the chordate lineage is limited by the lack of neuroethological studies leveraging our closest invertebrate relatives.

Here we combine high-throughput video acquisition with pharmacological perturbations of bioamine signaling to systematically reveal the global structure of the motor behavioral repertoire in the *Ciona intestinalis* larvae. Most of *Ciona’s* postural variance can be captured by six basic shapes, which we term “Eigencionas”. Motif analysis of postural time series further demonstrated that bioamines influence numerous stereotyped behavioral maneuvers including “startle-like” and “beat-and-glide”. Employing computational modeling of swimming dynamics and spatio-temporal embedding of postural features revealed that subtle behavioural differences are generated at the levels of motor modules and the transitions between, both modulated by bioamines. Finally, we show that flexible motor module usage gives rise to diverse behaviors in response to different light stimuli.

## Introduction

A primary function of animal nervous system is to transform sensory input into a sequence of actions known as behavioral output. Thus, the overarching motive of neurobiology research is to delineate the functional makeup and mechanistic basis of these behavioral outputs. Major progress has relied on the development of experimental tools and analysis methods that permit real time measurements and quantitative characterization of behavior (reviewed in^1, 2, 3, 4, 5^). Among the various natural animal behaviors, locomotion forms an integral part of nervous system function. Researchers in the field have been able to employ the aforementioned modern technologies to define motor actions as a function of their natural stereotyped elements, known as behavioral ‘modules’, ‘motifs’, ‘syllables’, or ‘primitives’^1, 6, 7, 8, 9, 10^, where these basic building blocks of motor behavior operate under organizational and hierarchical rules that bear similarities to phonological and syntactical rules that govern language. Modern systems neuroscience approaches have greatly facilitated the investigation of vertebrate motor modules^11, 12, 13^, which in invertebrates are even more likely to be interrogated with high sensitivity and precision, largely due to the latter’s smaller nervous systems^14, 15, 16^. In addition, the next generation of neuroscience discovery capitalizes on developing and studying new non-traditional model species to reveal not only common principles, but also differences in behavioral organization across the tree of life as well as within important clades^17, 18^. Consequently, there is an urgent requirement for expanding neuroethological studies to additional organisms occupying key phylogenetic positions.

Invertebrate chordates belonging to the phylum Chordata are obvious candidates for neuroethological analysis since they are close relatives of vertebrates and may provide important insight into the evolution of chordate nervous systems. While the importance of studying invertebrate chordates has been recognized in the field of evo-devo, as evident from an explosion of evolutionary, genomic, and developmental studies primarily in three organisms: the cephalochordate Amphioxus and the tunicates *Ciona intestinalis* and *Oikopleura dioica*, these organisms have yet to be leveraged in the context of neuroscience. Understanding their nervous system functions and behavioral repertoire will provide insights into the conservation and diversity of locomotory circuits, and how these relate to the evolution of the diverse modes of locomotor behavior^19^.

Recent publication of the *Ciona intenstinalis* larval connectome^20, 21^, single-cell transcriptomes of the larval nervous system^22, 23^ and establishment of *in vivo* functional imaging^24, 25^ have made *Ciona* a promising target for functionally dissecting a small invertebrate nervous system at a systems level.

However, a major hindrance to employing Ciona larvae for systems neuroscience is the absence of a behavioral platform that can measure phenotypes in an extensive and intensive manner^26^, which is especially crucial for the analysis of locomotion due to its sensitivity to both neurogenetic^27, 28^ and neuropharmacological perturbations^29, 30, 31, 32, 33^.

In this work, we address this knowledge gap by using machine vision to track, skeletonize and extract postural features from thousands of larvae swimming both spontaneously and under light stimulation. We additionally combine wild type swimming behavioral analysis with a small scale pharmacobehavioral screen that targets bioamine signalling, a key regulator of the biophysical properties of neurons, synapses^29, 34, 35^ and behavior^29, 36^. Using dimensionality reduction, we derive lower dimensional representations of body postures, which we term “Eigencionas”, With these we are able to explain the majority of postural variance in the Ciona lavae. We also combine three state of the art approaches: Motif identification, Hidden Markov Model (HMM) and spatio-temporal embedding to quantitatively define *Ciona* larval behavioral dynamics and thereby uncover the perturbation-sensitive modulation effects exerted on them by bioamine neuromodulators, which are likely left undetected with less sensitive methodologies.

## Results

### Parametrization of Ciona using interpretable features

Using five inexpensive USB microscope-based tracking setups we recorded high-resolution videos of 1463 individually and freely swimming *C. intestinalis* larvae of which 694 were wild type larvae swimming in Artificial Sea Water (ASW) and 769 were incubated with drugs that target different neuromodulators (Table S1, Table S2), (Fig. 1A).

**Fig. 1.**
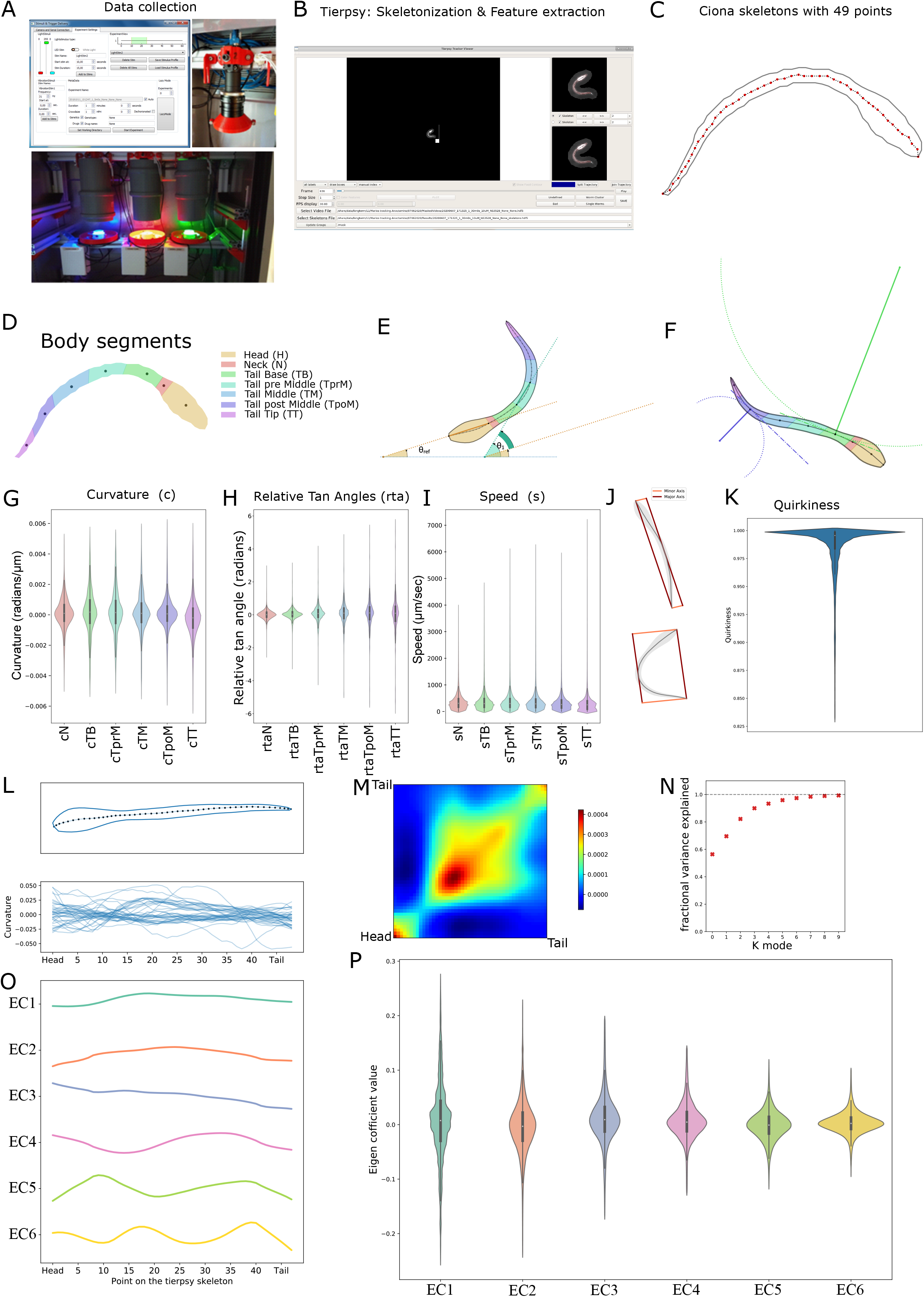
(A) Typical organization of a video acquisition setup. Screenshot of the software GUI we use to control the trackers. (B) We used Tierpsy analysis software in order to segment swimming Ciona larvae. Screenshot from Tierpsy Tracker viewer(*37*) showing a segmented and tracked Ciona. (C) Ciona skeletons generated using Tierpsy are defined by 49 equally spaced points on their midlines. (D) Skeletons were divided into seven body segments along the length of the animal. (E) Definition of angles: Θ1 refers to the tangent angle of a particular segment, θref refers to the tangent angle for the head segment (with respect to the horizontal axis). The difference θ1-θref is defined as the relative tangent angle for the particular segment. (F) Curvature definition visualization. Curvature is defined as inversely proportional to the radius of the osculating circle at a given point on the skeleton. Green shows lower curvature and blue shows the centroid of a segment with a higher curvature. (G-I) Distribution of curvature, relative tangent angle and speed values for each body segment in wild type larvae (n=694 larvae). (J) Quirkiness is defined as the ratio of the major and minor axes of the body as illustrated. (K) Distribution of the quirkiness for wild type larvae (n=694 larvae). (L) A skeleton with the contour and 49 points are shown as an example. Curvature values along the 49 points for a set of randomly sampled skeletons shows variation in the skeleton postures. (M) Covariance matrix calculated from curvature values of a subset of wild type skeletons. The smooth structure of the covariance matrix indicates that postures can be represented using a small number of eigen-cionas. (n=231 larvae) (N) Six eigen-cionas are sufficient to explain 97% of the variance in the curvature. (O) Visualization of the top 6 eigen-cionas obtained by an eigen decomposition of the covariance matrix, shown in descending order of the fraction of the variance explained. X-axis refers to points along the skeleton. (P) Distribution of eigen coefficient values for wild type larvae (n=694 larvae). For statistical analysis we first tested for data normality using Shapiro-Wilk test (α=0.05). To compare between different body segments, we used the Wilcoxon signed-rank test (α=0.05) (Tables S3-S6).

We then utilized the Tierpsy software^37^ to extract, in a high-throughput manner, 49 two dimensional positional coordinates of each of the contours and midlines of each Ciona larva, from our videos. This allowed us to approximate all the larvae via the 49 positional coordinates of their midlines (skeletons) during our downstream analysis (Fig. 1B,C).

Next, we grouped the 49 points identified by Tierpsy into 7 distinct segments ranging from head to tail (Fig. 1D and see Methods). We are more interested in the six segments from neck (N) to tail tip (TT), given our assumption that the head segment is rigid. We then defined relative tangent angles for each of these six segments relative to the head segment (Fig. 1E). The relative tangent angles provide a measure of the orientation of each of the segments with respect to the head segment. Curvature values, on the other hand, give a quantitative measure of the local bend at the middle of each of the segments, which is independent of the overall shape of the larva (Fig. 1F). This difference can be seen in the violin plots of Fig. 1 where the wild type larvae curvature in each of the segments have a similar range of values (Fig. 1G) whereas for the relative tangent angles, the range (variance or spread) of values become wider as we move away from the neck segment (Fig. 1 H). The highest mean segment speeds are seen in the neck region (sN), while the lowest mean speeds are observed at the tail post middle (sTpoM) and tail tip (sTT) segments(Fig. 1I).

We also calculated quirkiness values which give a scalar measure of eccentricity. A quirkiness value of 1 would mean that the skeleton has a perfect straight-line shape and quirkiness values closer to zero would indicate a skeleton where the bounding box (Fig. 1J) is nearly a square that encloses a highly curved animal. The quirkiness distribution of the wild type dataset is in line with our empirical observation that the *Ciona* larvae, while stationary or swimming, primarily maintain a relatively straight body posture where exaggerated tail bends are rare. These are reflected in the lower tail of the violin plot (Fig. 1K).

We further investigated if a lower dimensional feature space exists to capture the rich and diverse postural features exhibited by the *Ciona* larvae, as represented by a randomly sampled set of curvature values (Fig. 1L). While features like curvature and relative tangent angles could describe postures very accurately, the richness in these descriptions comes at a cost of very high dimensionality. Our aim was to obtain a simpler representation which describes the wide range of postures without losing significant information. We initially examined if there are any dependencies or relationships between the 49 points on the skeleton by looking at the covariance matrix of curvature values from 231 experiments, amounting to over 2 × 10^6^ images, or skeletons (Fig. 1M). As expected this matrix indicates a strong correlation between adjacent points, indicating that the 49 points do not move independently of each other. We confirmed this by performing an eigenvalue decomposition of the covariance matrix and found that 97% of the variance observed in the curvature data can be explained by 6 eigenvectors (Fig. 1N). For any given frame in the video, the curvatures of the skeleton can be approximated as a linear combination of these 6 ‘Eigenciona’ vectors: EC1 to EC6 (Fig. 1O). Hence the coefficients or scores of these 6 eigenvectors were used for further analysis as a simpler but nearly accurate description of skeleton postures (Fig. 1P).

Following the calculation of the biophysical features as described above, we looked at the statistical differences in each of them across the wild type and drug-treated experimental groups. We employed different metrics and visualizations for this purpose. First, we used Standardized Median Difference (SMD)^38^ to compare the distributions of features obtained from the different drug datasets with our wild type dataset (Fig. 2 A). The SMD values demonstrate that for the segments speeds’ the largest increase is observed upon treatment with phentolamine, which is an α-adrenergic antagonist and raclopride a D2 dopamine receptor antagonist, while the largest decrease is observed in animals treated with imipramine, a potent serotonin reuptake inhibitor (Fig. 2A, Table S7). For quirkiness, Chlorpromazine, an antipsychotic drug, has the lowest SMD value; while quinpirole, a selective D2/D3 receptor antagonist, exhibits the highest SMD value. Multiple drugs showed an overall increase in body curvature and relative tangent angles across most body segments. Paroxetine, a serotonin reuptake inhibitor (SSRI) and Phentolamine showed the biggest increase relative to wild type. In contrast both raclopride and quinpirole showed substantially decreased values for these features (Fig. 2A, Table S7). These trends are also observed in the visualizations using split violin plots of the distributions (l Fig. S1 A-C, Fig. S2 A,B).

**Fig. 2.**
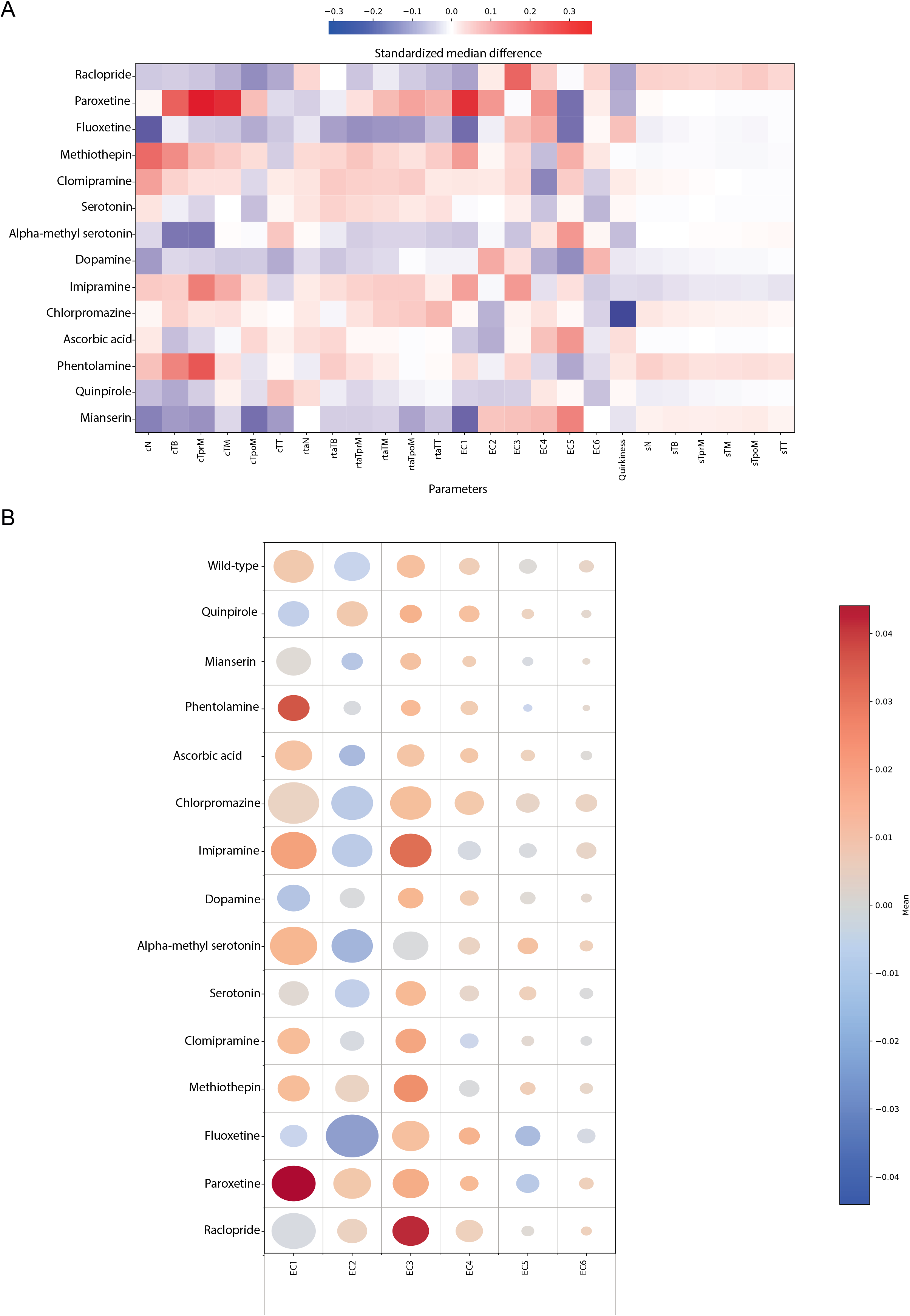
(A) Heatmap showing Standardized Median difference (SMD) of 25 features, calculated for drug treated larvae relative to wild type larvae. Note that SMD values for Dopamine are calculated relative to an Ascorbic Acid solution which was used as a solvent for Dopamine (SMD values shown in Table S7) (B) Bubble grid chart showing effects of drug treatments in the use of eigen Cionas by swimming larvae, as quantified by the eigen coefficients. Colour of circles indicates the mean value of the eigen coefficient features as indicated by the colorbar, while their radius indicate the variance. The number of animals and video frames contributing to this figure are indicated in Table S1-S2.

Then, to examine the differences in the eigen coefficient features, we present a bubble grid chart (Fig.2 B) where the hue and radius of the circles respectively represents the mean and the standard deviation of the distribution. For EC1, phentolamine and paroxetine have a large positive mean for EC1, suggesting that the EC1 component largely represents the shapes observed in larvae treated with these drugs. Conversely, for EC2, fluoxetine has a large negative mean with high standard deviation. Similar to EC1 and in contrast to EC2, EC3 exhibits a very strong positive trend across most drug treatments. Notably, the EC3 component heavily contributes to the skeleton postures in larvae treated with imipramine and raclopride. The remaining eigen coefficients, EC4 to EC6, show comparatively modest changes in mean values compared to control, except for EC5 in larvae treated with the SSRIs fluoxetine and paroxetine that exhibit a strong reduction in the mean value of the distribution. Overall, eigen coefficients are good descriptors of skeleton posture; and at least EC1 to EC3 are shown to be upregulated and downregulated in a drug-specific manner in our pharmacobehavioral screen

### Ciona locomotion is rich in behavioral motifs across timescales

To measure the stereotypy and reveal the modular structure in the motor repertoire of *Ciona*, we used a multitude of state-of-the-art analytical methods including motif discovery, Hidden Markov Models (HMM) and spatio-temporal embedding into a lower dimensional space. The first approach we took was that of recurring motif discovery. Recurring motifs indicate that some information is conserved for a system to produce the same output at least twice, in our case a repeated behavioral action or state. To perform automated behavioral motif detection, we decided to use matrix profile analysis^39, 40, 41^, since it is robust, scalable, computationally efficient, largely parameter free, and capable of identifying both recurring motives and anomalies in the data. For our analysis we have employed the curvatures of the seven body segment midpoints of the *Ciona* larvae. This results in seven-dimensional time-series which we used to calculate matrix profiles and search for recurring motifs over two time-windows: 1 second (30 frames) and 5 seconds (150 frames) in order to capture both short (spontaneous) and long (sustained) behaviors that repeat over time (Fig. 3 A). We identified the motifs over the two time-windows which resulted in two datasets consisting of a set of 87569 motifs over 1 second, and a set of 18776 motifs over 5 seconds. In the 1 second time-window we find repeating motifs that correspond to larvae performing different swimming manoeuvres including clockwise (CW) or counter-clockwise (CCW) turns, straight runs, twitching, rapid accelerations, or decelerations, beat-and-glide, as well as startle-like escape actions (Fig. 3B). In the 5 second time-window we found a lot of CW and CCW spiral swimming, circle swimming, straight runs which conclude in different ways: rapid holt, swim in a small circle or perform a spiral swim. Other motifs include drifting, persistent unidirectional tail flicking and again startle-like escape actions that take place over a longer time-window (Fig.3C).

**Fig. 3.**
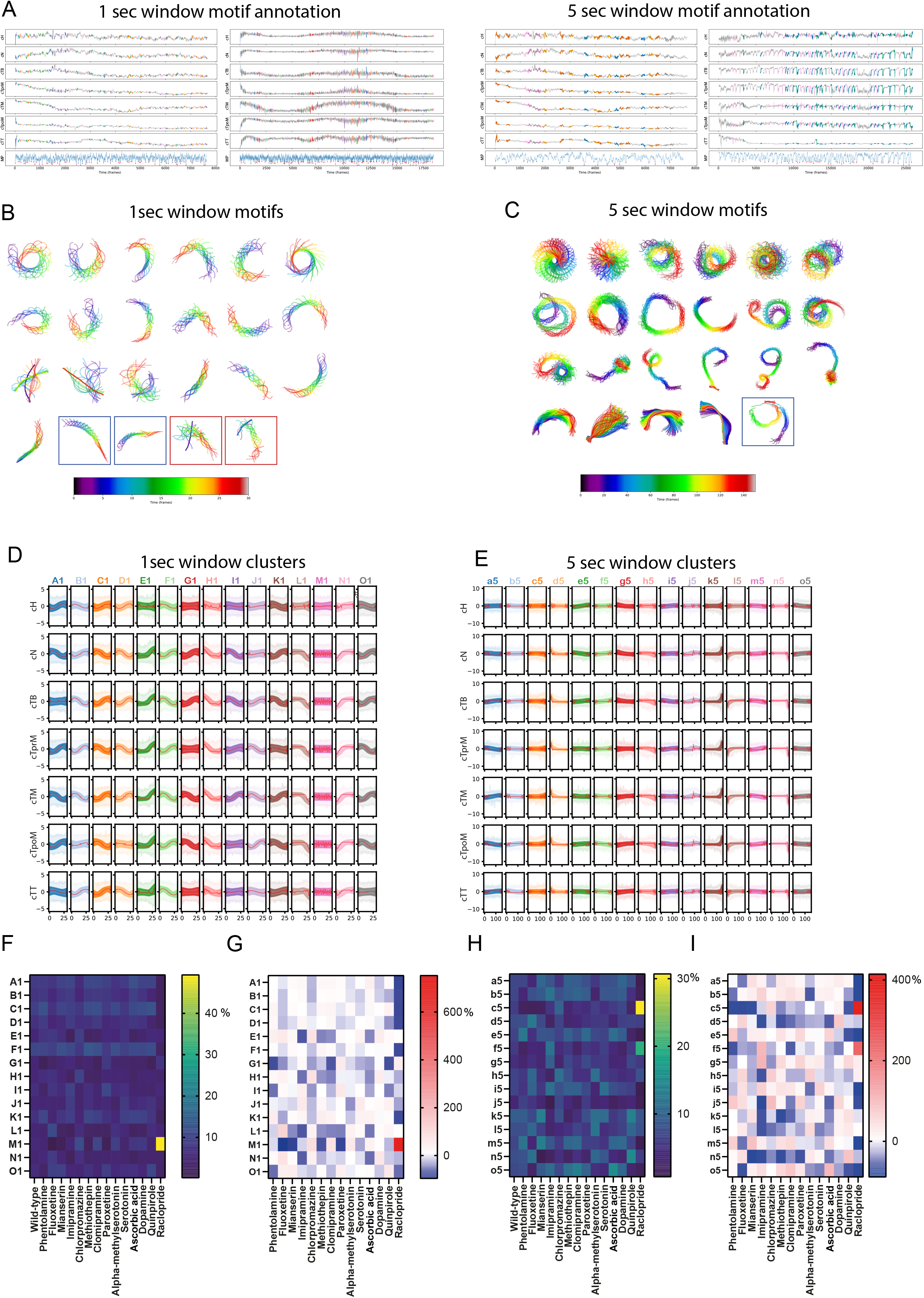
Four representative cases of larval swimming analyzed using matrix profiling. Curvatures of 7 body segments were used as the input. The matrix profile for each animal is shown as an additional row at the bottom of each plot (labelled as MP). Each dip marked with a red dot indicates the onset of a motif that recurs in the dataset. Each motif is colour coded according to the cluster it belongs to. (B) Representative examples of motifs that are enriched in the 1 second time-window. Color-coded to show time progression (0→30frames, i.e. blue→red). Enclosed in the blue boxes are example beat- and-glide motifs and in red boxes startle response motifs. (C) Representative examples of motifs that are enriched in the 5 second time-window. Color-coded to show temporal progression (0→150frames, i.e. blue→red). Enclosed in a blue box is beat-and-glide motif. (D) Timeseries clustering of motifs for 1 second time-window and (E) for 5 seconds time-window. In the graphs each column corresponds to one motif cluster while each row corresponds to one of the seven body segment curvatures. Each cluster was assigned a colour and a letter. Red line in all clusters indicates cluster centre and the variance is shown as shaded lines. Use of capital letters in (D) and small letters in (E) is done to indicate that these are not the same clusters. (F) Heatmap visualization of the 1second time-window motif clusters representation (values in % can be found in Table S14). (G) Heatmap visualization of the percentage fold changes relative to control for the data shown in panel F(values in % can be found in Table S15). (H) Heatmap visualization of the 5 seconds time-window motif clusters for different drugs (values in % can be found in Table S16). (I) Heatmap visualization of the percentage fold changes relative to control for the data shown in panel H (values in % can be found in Table S17). Dopamine values are compared relative to Ascorbic Acid.

We then asked whether there are differences between these motifs. Given that our two motif datasets are practically speaking large sets of short seven-dimensional timeseries, we performed timeseries clustering using k-means clustering (TimeseriesKMeans) to identify major groups to classify our motifs into. We determined the optimal number of clusters to be 15 for each of the time-windows (Fig. 3 C, D) using the Elbow method, which identifies the point at which adding more clusters to a model decreases the inertia of the model considerably less. From these clusters, the following involved actively swimming larvae (1sec: “H1”, “L1”, “M1”, N1”; 5sec: “c5”, “d5”, “j5”, “l5”) while other clusters (1sec: “A1”, “D1”, “E1”, “G1”; 5sec: “f5”, “g5”, “h5”, “k5”, “n5”) represent moderately active and (1sec: “B1”, “C1”, “F1”, “I1”, “J1”, “K1”, “O1”; 5sec: “a5”, “b5”, “e5”, “I5”, “m5”, “o5”) represent dwelling larvae. Random sampling of single frame skeletons as well as 1 and 5 seconds long sequences of skeletons revealed behaviors that set apart the different clusters (Fig. 3D,E, Fig. S3 A-D Movies S1-S30). For example, in the active clusters, “M1” we find motifs corresponding to larvae performing sharp CW or CCW turns with high curvature tail beats, while clusters “L1 ” and “N1 ” are enriched in larvae showing startle-like behaviors and unidirectional tail flicking characterised by asymmetric tail beats. We were also able to identify biologically interesting clusters within the 5 seconds time-window, such as active cluster “c5”, which is enriched in CW and CCW circular and spiral swimming motifs as well as straight runs concluding with a circular or spiral manoeuvre (Fig. S3 D, Movie S18). Cluster “d5” is enriched in short range swimming and late onset escape manoeuvres (Fig. S3D, Movie S19). Interestingly, while cluster “l5” was enriched in motifs where most of the swimming activity occurred in the first 3 seconds (Fig. S3 D, Movie S27) clusters “k5” and “n5” were enriched in motifs where the majority of the swimming activity occurred within the 2 last seconds of the 5sec time-window (Fig. S3 D, Movies S26, S29).

For the 1 second time-window behavioral motifs belonging to clusters “A1” to “D1”, “F1” and “K1” are the most frequently identified across the wild type data (Fig. 3 F; Fig. S3 E; Table S14). This is consistent with our empirical observations that *Ciona* larvae spend a considerable time slowly swimming or staying idle. We then asked how the different drugs we applied have affected the motif cluster distribution. Raclopride treatment results in a strong reduction in the representation of Clusters “A1” to “D1”, (Fig. 3 F, G; Table S14, S15). Suggesting that dopamine signalling is important for moderately active swimming and dwelling behaviors. Clusters “E1” and “H1” that represent moderately active and active larvae respectively, show a similar profile in response to the drugs used in this study (Fig. 3 F, G; Fig S3 E, F; Table S14, S15), indicating that the behavioral motifs included in these clusters may be generated by a common underlying cellular and/or molecular mechanism. In sharp contrast, the remaining inactive clusters “F1”, “I1”, “J1” and “O1” have little in common in their response across the drug treatments (Fig. 3 F, G; Fig S3 E, F; Table S14, S15). Amongst the active clusters “L1”, “M1” and “N1” are strongly modulated. Raclopride results in a very strong upregulation of cluster “M1” indicating that normally dopamine signalling serves to suppress sharp CW or CCW turns. Interestingly, Fluoxetine shows very strong phenotypes across most of the 1 second active clusters where it strongly upregulates clusters “L1” and “N1”, while substantially downregulating cluster “M1” (Fig. 3 F, G; Fig S3 E, F; Table S14, S15 This suggests that serotonin signalling likely suppresses startle-like behaviors and unidirectional tail flicking. Taken together the results from raclopride and fluoxetine suggest that dopamine and serotonin signalling exert opposite effects on CW and CCW turns (Cluster M1).

In the 5 seconds time-window, we observed a uniform representation of most clusters in wild type data, except for the active cluster “j5” (Fig. 3 H; Fig. S3 F; Table S16). What can be readily appreciated is that most of the motif clusters in the 5 seconds time-window are more strongly regulated by the drug treatments in comparison to the 1 second time-window. The drug with the most dramatic changes in cluster representation is Raclopride followed by Imipramine (Fig. 3 I; Fig. S3 F; Table S17). Raclopride treatment strongly increased the usage of CW and CCW spiral and circular swimming cluster “c5” as well a as the moderate activity cluster “f5”, while strongly suppressing various low swimming activity clusters (“a5”, “b5”, “m5”, “o5”). Imipramine on the other hand showed a very strong redistribution of cluster usage within the moderately active ones downregulating clusters “k5” and “n5”, which are enriched in motifs that show a late onset of swimming manoeuvres and upregulating two clusters where twitching (“f5”) and spiral swimming (“g5”) manoeuvres unfold throughout the 5 seconds time-window. Interestingly, phentolamine, fluoxetine and mianserin treatments strongly suppressed the use of CW and CWW swimming cluster “c5” suggesting that serotonin and noradrenaline promote turns in swimming larvae.

### Ciona motor behavior can be modelled in terms of states and transitions

Having demonstrated that motif identification using matrix profiling is a potent method for identifying the basic behavioral building blocks of *Ciona* behavior, we next sought to expand our work by performing a systematic analysis of the organization of the behaviors that can be performed by the *Ciona* larvae. With an underlying assumption that behavior is modular and can occur across multiple timescales, we modelled our behavioral data with a simple Gaussian Hidden Markov model (G-HMM). HMM provides a dynamical framework to identify the distinct behavioral modules that repeat over time at different time scales^8, 42, 43, 44, 45^. We implemented a 10 state HMM to model larval swimming across different experimental conditions (see Methods). We analyzed the means and standard deviations of the features for each of the states (Fig. 4 A) to show that the states β, γ η and κ have one or more input features with a distinctively higher standard deviation. This indicates that the larvae, while in any of these states, take a range of feature eigen components and/or quirkiness, suggesting variation in postures that can be inferred as a result of active swimming. We then produced animations of skeleton movements in the arena for each of the states (Fig. S4; Movies S31-S40). The animations agreed with our inference that states β, γ η and κ correspond to active swimming in the arena.

**Fig. 4.**
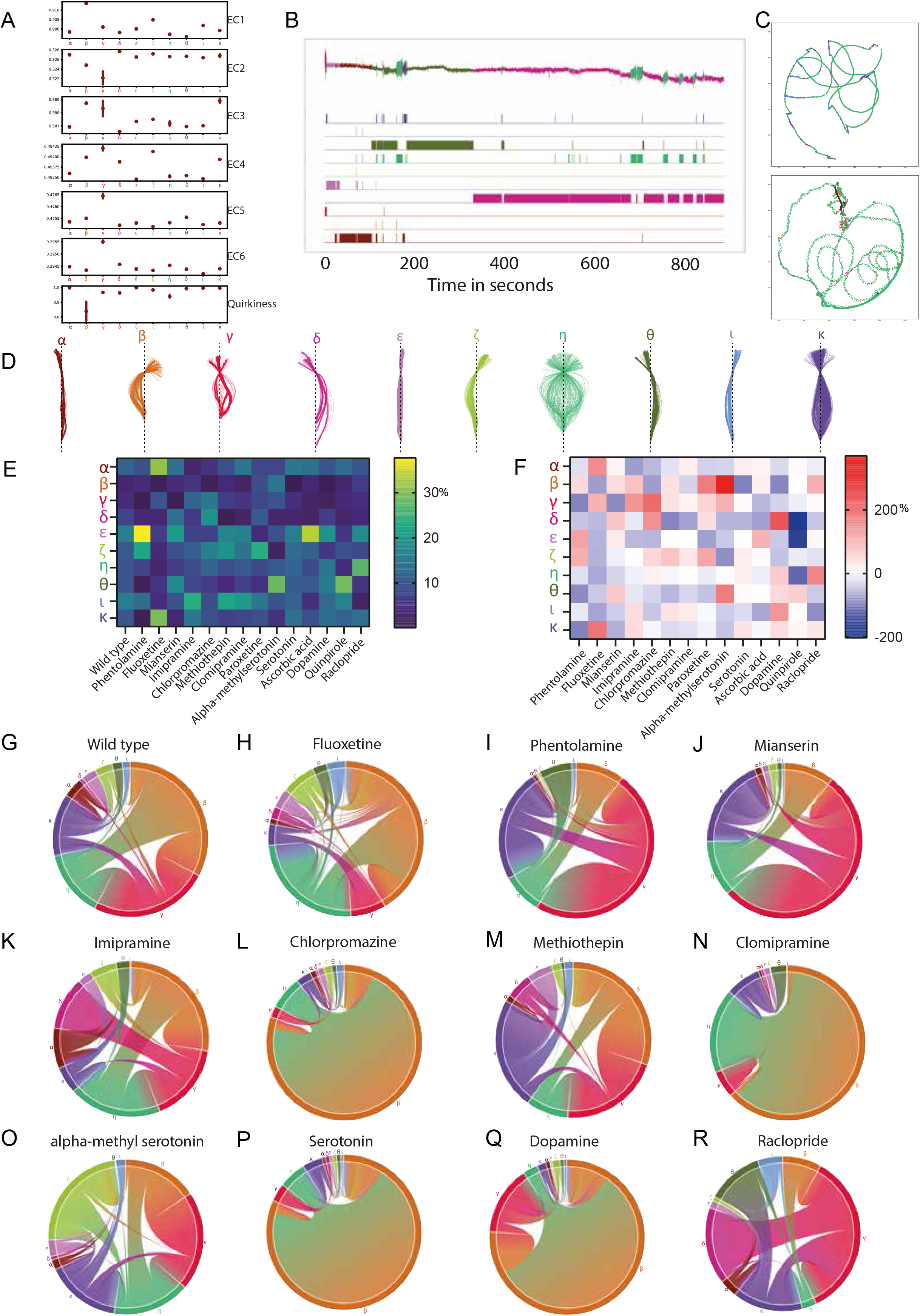
(A) Plots summarizing the observation probability distributions defined by the HMM model. For each of the states (horizontal axes), the mean of Gaussian distributions of each of the seven input features is plotted along the vertical axes. The variance of the distributions is indicated by the error bars. (B) HMM segments and clusters a time series into modules by identifying the underlying state for each timepoint in the series. On the top eigen coefficient EC1 (a time series) of a larva swimming is annotated with the different HMM states (uniquely color coded) identified in our analysis. (C) Two example tracks of the neck point of larvae in the arena coloured according to the behavioral state identified by HMM. (D) Postures /skeletons were randomly sampled 90 skeletons from the dataset for each of the ten different HMM states, aligned such that the neck points coincide and are collinear with tail-ends on a vertical line. (E) Heatmap visualization of the effects of drug treatments on the HMM-derived behavioral states (values in % can be found in Table S18) (F) Heatmap visualization of percentage fold changes relative to control for the data shown in panel E (values in % can be found in Table S19). Dopamine values are compared relative to Ascorbic Acid. (G-R) Chord diagrams showing HMM derived behavioral state transitions for wild type and drug treated larvae. Chord diagrams are presented in such a way that transition to self (i.e., persistence of states) are not shown so that the transition probabilities across states can be highlighted. Self-transitions and transitions with probabilities less than 0.001 are ignored.

We also assessed if our model can predict underlying states consistently across experiments and datasets. For this purpose, we labelled time series of input features by the inferred state obtained from the model prediction Fig. 4B. The model suggests that the larvae exhibit intermittent locomotion with bursts of swimming and substantial periods of dwelling. Using the inferred HMM states, we visualized the trajectories of the neck point along the arena for different experiments (Fig. 4 C; Fig. S5 S-V). Importantly, these visualizations highlight a new behavior that resembles the beat-and-glide behavior observed in zebrafish larvae^46^. This is an intermittent form of swimming defined by tail beating followed by gliding during which the tail remains relatively motionless either straight or with a small amount of curvature. State “η”, which is one of the actively swimming states, is the dominant ‘beating’ state which is followed by one of the multiple gliding states “α”, “δ”, “ζ”, “ι”. This set of skeletons shows that the range of postures within each of the states are highly consistent. For example, states “α”, “δ”, “ε”, “ζ”, “θ”,and “ι” represent different idle phases each with unique skeleton shapes. For states “α” and “ε”, the sampled skeletons are close to a straight line, which reaffirms the observation in Fig. 4 A, where they have the highest quirkiness values with negligible variance suggesting that these states do not correspond to active swimming. Similarly, another trend revealed by our analysis is that skeletons belonging to states “β”, “ζ” and “ι” are curved towards the left side of the vertical axis in Fig. 4 D and have higher values of EC1 feature. The wide range of skeleton shapes in state “η” (Fig. 4D) agrees with the fact that active swimming and exploration of the arena would require the larva to take different postures at various stages within a single cycle of swimming motion. This is also confirmed by lower mean values of quirkiness for state η (Fig. 4 A).

Using the results from HMM, we examined how the distribution of distinct behavioral modules varies across the different drug treatments by calculating the percentage of representation of each of the states for different drug datasets. For example, as shown in Fig. 4E, F (see also Table S18, S19), we demonstrate that in Fluoxetine, states “α” and “κ” are overrepresented in comparison to the wild type. On the other hand, certain states are heavily underrepresented or entirely absent when the larvae are treated with certain drugs. For example, state “β” is strongly underrepresented in Fluoxetine, Chlorpromazine, Clomipramine a serotonin-noradrenaline reuptake inhibitor (SNRI), exogenous Serotonin and Quinpirole (Fig. 4 E; Fig. S5 A, B; Table S18, S19). Interestingly, the serotonin receptor agonist alpha-methyl serotonin shows the strongest increase in the use of this active swimming state. The active state “γ” has a very low representation in drug treatments with Phentolamine and Mianserin. This suggests that noradrenaline and serotonin are important signalling molecules for promoting active behavioral states. Compared to wild type and other drug treatments, Imipramine and Raclopride have an overall increase in the representation of the actively swimming states and a significantly lower representation of the idle states “α” and “ε” where the animals adopt a nearly straight skeleton (Fig. 4 E; Fig. S5 A, B; Table S18, S19).

Similarly, the transition probabilities obtained from an HMM allows us to look at how different drugs affect the transitions from one behavioral state to another. This is visualized using chord diagrams (Fig. 4 G-R; Fig. S5 A-C) and Markov transition graphs (Fig. S5 D-R). We found that in wild type animals, transitions between states “β”, “γ” and “η” were the most prevalent, forming a core transition module. Beyond this core module active state “κ” acts as a ‘transit hub’ for most behavioral state sequences that occur at a lower frequency. Importantly, state transitions do not occur in an all-to-all fashion (Fig. 4G; Fig. S5D). For example, state “δ” interacts exclusively with state “γ” (Fig. S5 D). Our wild type data suggest that certain states can interface with multiple other states, while other states may be more exclusive in their interactions. Remarkably, we found that the number of state transitions and the plausible pairwise combinations are sensitive to pharmacological treatment. Our chord diagrams and transition graphs reveal that Clomipramine and Imipramine reduce the transitions between behavioral states (Fig. 4K,N; Fig. S5K, L). Conversely, Fluoxetine and Phentolamine result in an increase of transitions amongst behavioral states (Fig. 4H,I; Fig. S5E, J). The “β”←→“γ”←→“η” module is preserved across all pharmacological treatments and the same holds true for a number of state transitions (e.g., “α”κ-”, “ε”-”κ”, “η”-”θ”), however other transitions show drug dependent ‘plasticity’ (e.g., “γ”-”δ”, “β”-”ι”, “α”-”ε”). For example, state “δ” is solely interacting with the active state “γ” in the wild type dataset (Fig. S5E). Some drug treatments such as Imipramine strengthen the transition between these two states (Fig. 4F; S5 L). However, in alpha-methylserotonin treated animals “δ” interacts with state “θ” in (Fig. S5 I) or “η” in the case of clomipramine treatment (Fig. S5 N). Notably, we found that “δ” is not limited to interacting with one state but it can interact with multiple transition partners as observed in a subset of drug treatments (Fig. S5 F, G, J,K, N-P,R, S). Additionally, inferences can be made by combining the information from the percentage representations of states and the chord diagrams. For example, state “α” does occur at a very high percentage in larvae treated with fluoxetine, however, it can be seen from the chord diagram that the transition from “α” state occurs very rarely in these larvae, which indicates that they tend to stay idle and locked into state “α” for a long time.

### Low dimensional spatio-temporal embedding identifies stereotyped actions in swimming larvae

While we have made assumptions that behavior is organized with units of repeated motifs (matrix profiling) or modules with Markov transitions (HMM), we wanted to additionally adopt a complementary approach aiming to uncover new structures in our behavioral data. For this purpose, we employed an approach similar to Berman et al^7^ to reveal stereotyped behaviors exhibited by the *Ciona* larvae.

For this approach, we used as the input the 6 eigen coefficient features from a subset of the wild type data where it was sampled based on speed (Fig.5A). The sampling was performed to ensure that the actively swimming epoch is well represented. First, to encode the temporal information such that clustering is applicable, we created a 180 dimensional feature set by computing the wavelet transformation of 6 eigen features at 30 different frequencies or scales (Fig.5 B). Wavelet transformation allows us to create for each timepoint a feature that has information about its surrounding timepoints built into it, thereby creating a feature set where temporal information is preserved.

**Fig. 5.**
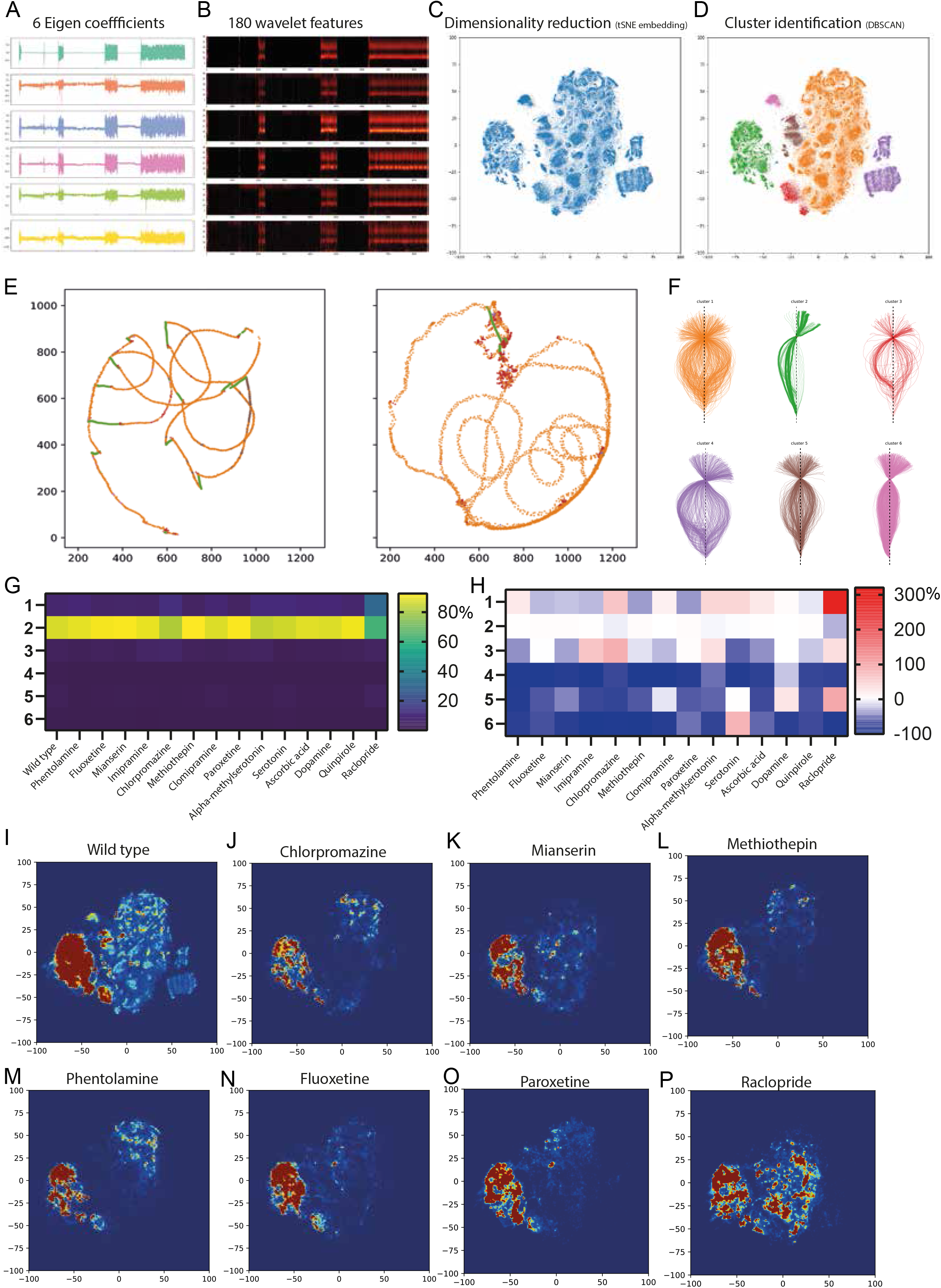
(A) We used the six eigen coefficients we previously obtained to define a six-dimensional input feature space. (B) We computed wavelet transforms of these features over 30 frequencies for the entire control dataset. Wavelet transform of the 6D eigen time series of panel A shown. (C) Dimensionality reduction was applied using tSNE to embed the 180-dimensional wavelet features into a 2D space of behavior. (D) We then used the DBSCAN algorithm to cluster into regions based on their density. (E) Examples of neck point tracks for larval swimming labelled according to cluster identity. (F) Examples of skeletons aligned to the neck for each of the clusters. (G) Heatmap visualization of the effects of drug treatments on cluster usage (values in % can be found in Table S20). (H) Heatmap representation of the percentage fold change relative to control for the data shown in panel [G] (values in % can be found in Table S21). In panels G and H Dopamine values are compared relative to Ascorbic Acid. (I-P) 2D tSNE embedding of wild type and different drugs. The color is showing the density with which the different clusters are occupied (blue being lower and red higher).

Next, to obtain a lower dimensional behavioral space from this feature set, we used t-distributed stochastic embedding (tSNE), which provides an embedding in which local structure is retained unlike many other dimensionality reduction techniques (Fig.5C). This embedding, which we can think of as the larval behavioral space, was then clustered using the DBSCAN algorithm (Fig. 5D). We identified 6 distinct clusters corresponding to different stereotyped behaviors (Fig. 5D). The DBSCAN, our algorithm also learned an additional outlier class which corresponded to less than 0.001% of the data points where cluster assignment was ambiguous (shown in blue in Fig.5D).

Our results show that the clusters identified by this method are coherent across datasets. For example, cluster 1 consistently represents video frames where the larva is actively swimming and exploring the arena. This can be seen in the trajectories of the neck point across 2 different experiments where cluster 1 (orange) dominates the phases when there is movement across the arena (Fig. 5E). On the contrary, cluster 2 (green) represents phases where the larvae were gliding in the arena (Fig. 5E). By inspecting a number of animal trajectories we were able to identify instances of beat-and-glide behavior like those observed using the HMM method. We provide further confirmation that the clusters we identified are coherent by randomly sampling skeletons across experiments for different clusters and plotting them such that the neck point and the end point are aligned on a vertical axis (Fig. 5F). Here, cluster 2 has the least variations in the skeleton postures it represents. In addition, examining the trajectories and snippets of animations of skeleton movements uncovered that cluster 3 (red) represents sharp turns associated with transitions from a gliding phase (for example, cluster 2) to an actively swimming behavior (for example, cluster 1) (Fig. S6; Movies S41-S46). In addition, from a detailed inspection of trajectories of the skeleton in the arena, we have identified that clusters 5 (brown) and 6 (magenta) represent swimming with lower speeds compared to the higher speeds exhibited by clusters 1 (orange) and 4 (purple). A similar trend can be seen in Fig.5F where the former clusters have a narrower range of postures in comparison to a wider range of skeleton postures exhibited by the clusters 1 and 4.

Subsequently, we trained a kNN classifier on the clustering results, and used this classifier to assign cluster membership to datapoints from the drug-treated dataset. This approach lets us compare how larvae across different experimental groups utilized the behavioral space (Fig. 5G, Table S20). In comparison to wild type, the drug-treated larvae utilize to an even more limited extent the active swimming cluster 4 (purple) (Fig. 5 G, H; Fig. 6 I-P; Fig. S6 G-H; Table S20). Cluster 1 usage is strongly upregulated by Raclopride and to a lesser degree by Chlorpromazine, alpha-methyl serotonin and exogenous serotonin (Fig. 5 G, H; Fig. 6 J,P; Fig. S6 J,M; Table S20). In contrast the SSRIs paroxetine and fluoxetine and the two structurally similar antidepressants Mianserin and Methiothepin show a substantial reduction of Cluster 1 (Fig. 5 G, H; Fig. 6 K,L,N,O; Table S20). Sharp turns that occur when larvae transition from idle to swimming which correspond to cluster 3 are strongly downregulated by exogenous serotonin, while chlorpromazine treatment results in the strongest upregulation (Fig. 5 G, H; Fig. 6 J; Fig. S6 J; Table S20).

**Fig. 6.**
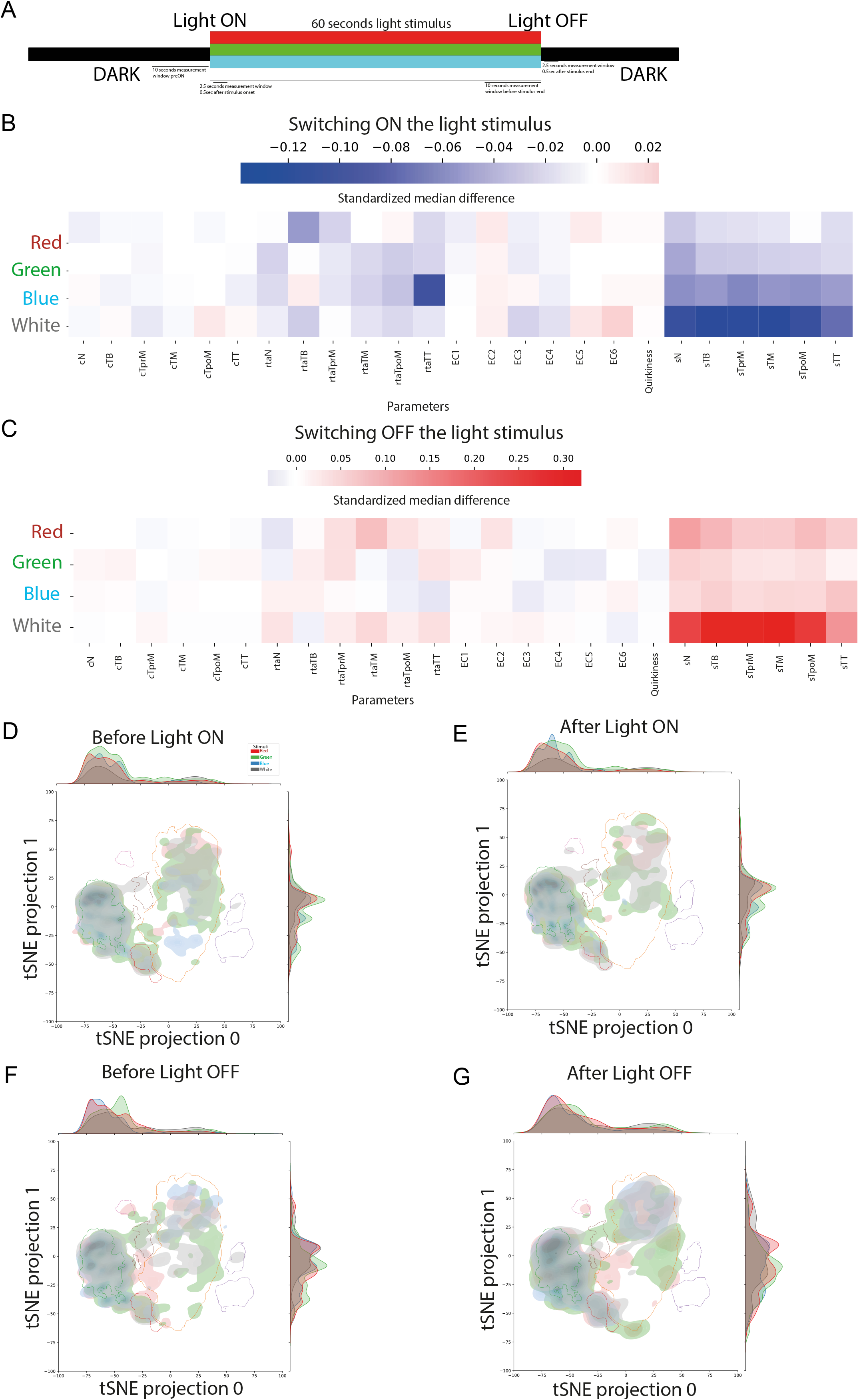
(A) Design of the light stimulus: A 60 second long window of stimuli is presented at the 30th second of the experiment. Different types of light stimuli are used, namely red, green, blue and white. We observe the changes at light ON event by comparing a 10 seconds window before the event (before Light ON) and a 2.5 seconds long window after 0.5 seconds of the onset of light ON (after light ON). Similarly, we define two windows before and after the light OFF event to study the changes in behavior when the animal exits the stimulus period. (Number of animals and frames used to generate this figure are indidcated in Table S1 & S2) (B) Effect of light ON event in the different biophysical features for each of the four stimulus types. The effect is measured in terms of the standardized median difference (SMD) in the feature values between the ‘before light ON’ and ‘after light ON’ intervals. (SMD values are shown in Table S22) (C) Effect of switching OFF the different light stimuli in terms of SMD in presented, similar to B (SMD values are shown in Table S23). (D-G) 2D density plots showing the change in the pattern of occupancy of the 2D behavioral space across different stimuli before and after the light ON and OFF events. Boundaries of the 6 DBSCAN clusters are also shown on the plots (2D density plots for each stimulus individually are provided in Fig. S10).

Finally, we mapped the transition probabilities between the behavioral clusters which allowed us to get insight into the organization of larval behavior. For example, we could infer that wild type larvae executing the beat-and-glide behavior have multiple intermediate cluster options to transition from a ‘beating’ phase (dominated by cluster1) to a ‘gliding’ phase (cluster 2). However, exit from the gliding phase to the beating phase (in this case primarily clusters 1 and 4) almost always occurs through cluster 3, which is characterized by an asymmetrical swimming movement (Fig. S7A, Movie S47). The 2→3 and 3→4 transitions are ‘resistant’ to almost all pharmacological perturbations, except for 3→4 in Chlorpromazine animals. However the 3→1 transition statistics are subject to modulation but several drugs affecting serotonin, noradrenaline and dopamine signalling (Fig. S7F, H,-K, N). Additionally, pharmacological treatments can establish new interactions between clusters (e.g. 5→4) (Fig. S7F,K). These illustrate that Ciona locomotion has a certain degree of hierarchy and organization with some of the transitions being sculpted by bioamines.

### Light stimuli modulate postural dynamics and behavioral space occupancy

In addition to pharmacologically inhibiting bioamine neurotransmission, we sought to address how the presentation and removal of sensory cues such as light affect larval swimming behaviors.

Previous work has shown that ascidian larvae exhibit a shadow (i.e. looming-object escape) behavior as well as positive and negative phototaxis^47, 48, 49, 50^. To probe these questions, we examined how motor behavior changes when the larvae enter and exit a shadow stimulus period. Our SMD feature values reveal that once the larva enters the stimulus period, for a 2.5 seconds time-window, the swimming speed decreases significantly when compared to a 10 seconds window defined before the stimulus period (Fig. 6A, B; Table S22). Removal of the stimulus gives rise to an opposite effect. A significant increase in speed is shown via the SMD speed values, in a 2.5 seconds window after the larva exits the stimulus period, when compared to a 10 seconds window during the presentation of stimulus period (Fig. 6A, C; Table S23). We show a similar trend for SMD relative tangent angle values (Fig. 6B & C).

We additionally demonstrate a difference in responses across different light stimuli types: red, green, blue and white. The differences in SMD speed values are significant for white light in both switching ON and switching OFF the stimulus (Fig.6B, C; Fig. S8, S9; Table S24-S31). Blue light has a significant effect while switching ON the light (Fig. 6B, Fig. S8) but a smaller effect when switching OFF the stimulus (Fig. 6C, Fig. S9). On the contrary, the effect of red light while switching OFF the stimulus is larger than during switching ON (Fig. 6B, C; Fig. S8, S9).

We then asked whether the presentation and removal of different color light stimuli influence the behavioral space explored by the larvae. As expected, the presentation of a light stimulus (ON) reduces the use of the active clusters in favor of the lower activity clusters 2 and 6 for white light stimulus (Fig. 6D, E; Fig. S10D, H; Table S32). However, there seems to be light color specific use of clusters. For example, in contrast to the white light ON period, during the Red light ON period slow swimming behaviors under cluster 6 are not used. Our data suggests that the larvae gradually adapt to the continuous presence of the light stimulus especially in response to green and blue light as is evident from comparing the tSNE plots immediately after the onset (Fig. 6E; Fig. S10 E, F) and prior to the end of the light stimuli (Fig. 6F; Fig. S10 I, J). Subsequent removal of the light stimulus causes the larvae to increase their swimming activities and thus reuse the higher activity clusters of the behavioral space (Fig. 6G, Fig. S10 M-P).

## Discussion

### Motion tracking and comprehensive feature extraction at high-throughput, resolution and reliability

In this work, we have characterized the feature-rich swimming behaviors of the protochordate *Ciona intestinalis* at an unprecedented level of detail. Previous behavioral studies from ours and other groups that investigated locomotion in Ciona^48, 51^, ^52, 53, 54, 55, 56, 57, 58, 59, 60^, Oikopleura and Amphioxus (reviewed in ^19^), have been challenged by the lack of advanced tracking and analysis methods, resulting in a limited quantitative characterization of the structural organization of the invertebrate chordate’s motor behaviour. Several of the papers on ascidians performed tracking through estimated centroid positions ^48, 51^, ^54, 56, 61^, with only a couple of exceptions having gone through the effort of tracing the larval body midlines without assistance from the machine vision and analysis tools available to us today ^59, 62^. Thus, a major technological advancement of this work is the utilization of the lengthwise midline of the larvae obtained using the Tierpsy software^37^, at high-throughput and reliability over the entire duration of the larval swimming videos. Through this approach, we have derived a dimensionality-reduced representation of *Ciona* body postures which we term “Eigencionas” that is more comprehensive than the discrete biophysical features such as segment speeds, curvatures and tangent angles; classically important parameters for describing motor behavior. Remarkably, we show that just six basic shapes (Eigencionas) can be combined in different proportions to reconstruct almost the entirety (~97%) of *Ciona* postures during swimming. The use of eigen vectors as lower dimensional representations of posture has been well established in *C. elegans*^9, 63, 64, 65^, Drosophila ^66, 67^ and zebrafish^68^. We believe that analogous to what has been done in these mainstream model organisms, Eigencionas will be widely employed in future behavioral analyses in *Ciona* and other ascidians.

### Motif analysis enables generation of a multiscale dynamic behavioral representation

Behavior is a highly dynamic phenomenon that entails changes to an animal’s posture over time. The realization that a major fraction of animal locomotion is low dimensional and stereotyped has sparked the development of multiple approaches to quantify stereotyped behavioral dynamics across several model organisms (reviewed in^3, 4, 5^). Our study is one of the few to the best of our knowledge to use multiple complimentary approaches to break down the behavioral dynamics of *Ciona* to reveal its modular structure.

Our first method searched for and extracted motifs from tracking data to generate a dynamic multi-scale representation of Ciona motor behaviour. Comprehensive inspection of the plethora of identified motifs has highlighted that existing manual approaches would most likely overlook motifs that represent subtle or apparently irregular yet repetitive behaviors. For example, other than motifs that represent known behaviors that have already been characterised by our group (e.g. ‘twitching’)^51^, the current work has exposed the previously underappreciated wealth of CW and CCW exploratory swimming manoeuvres, which may reflect the asymmetry of motor pathways in the Ciona larval connectome^20^. In addition, here we discover a novel startle-like manoeuvre, which may be associated with a pair of descending decussating neurons (ddNs) found in the motor ganglion. ddNs’ ultrastructure, network connectivity and synaptic connections have been elucidated in a recent study that has postulated that this neuron pair show network homology to vertebrate reticulospinal neurons and that their synaptic connectivity resembles that of the Mauthner cells ^21^, which underlie fish startle responses ^69, 70^. Indeed, we demonstrate in this work that serotonin suppresses startle-like behavioral manoeuvres. In line with this result, serotonin has been shown to regulate startle responses in zebrafish^71^. Our findings thus suggest that motor behaviour as well as the underlying molecular and cellular players are conserved across invertebrate chordates and vertebrates.

### Model fitting preserves temporal component to expose biologically meaningful motor modules and transitions

To further examine the modularity and transition structure in Ciona behavioral dynamics across diverse spatiotemporal scales, we have built a Hidden Markov Model (HMM). Using HMM, we have shown that *Ciona* larval locomotion can be decomposed into 10 distinct stereotyped locomotor states that can occur over a wide range of spatiotemporal scales. This framework provides the first probabilistic model (in terms of Gaussian distributions) for each of the distinct motor behavioral states of the *Ciona* larvae.

This richness of locomotor states comes as a surprise, since they are the output of a relatively simple nervous system equipped with a minimal number of neurons that make up the motor circuit of the larva^20, 21^. This work has identified different low activity states that are distinguishable by different resting postures adopted by the larvae and high activity states with distinctive swimming speeds and bending asymmetries. Interestingly, we found that most of the low-activity states exhibit unilateral (either left or right-handed) tail bending/flicking, while one active state shows unilateral tail bending. Sided flicking and swimming, are likely generated by asymmetries in sensory input to the motor ganglion of the larva as suggested by the Ciona larval wiring diagram ^20^. In addition, we have identified a new behavior that shows strong similarity with the beat-and-glide behavior that zebrafish larvae perform^46^, In zebrafish, there are suggestions that this behavior is at least in part by dopamine though this is still debated^72, 73^. Our findings show that both serotonin and dopamine regulate the transition statistics between the beating and gliding states, pointing to a more complicated modulation of the beat-and-glide behavior across species compared to what was previously thought.

Another key feature of HMM is the ability to describe the organization of behavior across time in terms of transition probabilities. The transition structure in wild type behavioral sequences revealed a core module composed of two asymmetric slow swimming states and an active symmetric swimming state that is dominant during exploratory behaviors. It also identified an active state (“κ”) that acts as a hub for the less frequent transitions that occur between states outside this core module. Importantly, the time spent by a larva in each behavioral state and the transitions between states appear to be selectively modulated by bioamines.

### Construction of a *Ciona* larval locomotor behavioral space

In parallel to employing the above methodologies, we have implemented a spatio-temporal mapping approach similarly to what has been used in mapping complex behavioral responses in *Drosophila* and mice^7, 74, 75^. Here we present six annotated clusters of different behaviors that segregated to the different regions of the map.

Interestingly, as with the HMM method, the beat-and-glide behavior is also identifiable using the spatio-temporal mapping approach. Due to the smaller number of clusters generated by the latter method, it has been more straightforward to visually infer the cluster usage and the key transitions between clusters that are required to generate this behavioral manoeuvre. That, the key clusters and transitions involved in the behavior were preserved under all pharmacological treatments suggests that these features are controlled by neuromodulators other than bioamines e.g. GABA and Glutamate. However, the transition statistics were demonstrated to be affected by dopamine and serotonin (which is in line with the HMM findings), indicating that bioamines likely perform the fine sculpting of this behavior,

In addition, using behavioral mammping we could visualize behavior as a trajectory across a manifold and examination of the dynamics of the same. In this behavioral space, spontaneously swimming *Ciona* larvae can ‘navigate’ between the six behavioral islands of stereotyped behaviors in defined manners. This approach has been particularly informative in the light stimuli experiments, where we demonstrate that the behavioral responses and adaptation observed in response to different light color stimuli do not result in global changes in the underlying spatio-temporal structure, but rather they arise from the selective use of modules and changes in the transition statistics. This suggests that the larval brain can alter the use of individual modules and the transition statistics to generate responses to novel situations (e.g., sudden presentation of a sensory stimulus). This is likely a conserved strategy amongst invertebrates and vertebrates used to produce complex behavioral actions in response to sensory cues^8, 76^.

### The proto-hypothalamic territory of *Ciona* may regulate behavioral switching using bioamines

Here we have shown that Ciona larvae are able to use a finite number of behavioral modules that can be assembled in different sequences to generate diverse behavioural output that is likely modulated by internal state changes during spontaneous swimming or in response to sensory cues, in our case different light stimuli. This strategy has been observed across vertebrates and invertebrates^6, 77^. We demonstrate here that bioamines act as behavioral switches that modulate the transition statistics and overall time spent in different forms of active exploratory swimming and locomotor periods defined by distinct forms of dwelling, gliding and slow swimming. This is likely an evolutionarily conserved function^36, 78^. Dopamine in wild type animals promotes dwelling, gliding and slow swimming states, a phenomenon that has been observed in a number of organisms including zebrafish and xenopus^73, 79^. On the other hand, serotonin and noradrenaline promote active exploratory swimming rich in CW and CCW turns. Notably, in mammals arousal and waking states are stimulated by serotonin and noradrenaline ^80^.

An obvious question that arises from our study is, which cells and anatomical structures in the tadpole brain use bioamines to modulate the composition and organization of the larval behavioral repertoire? The dopaminergic cells are composed of a single cell cluster called the coronet cells, which have been characterized molecularly and homologized to the vertebrate hypothalamus^52, 81, 82, 83, 84^. The same cells express the serotonin transporter (CiSERT)^52^, though the rate limiting enzyme in serotonin synthesis tryptophan hydroxylase (TPH), is expressed in the vicinity of the motor ganglion and tail muscles^85^. Given that the vertebrate hypothalamus is also capable of modulating behavioral states by the secretion of neuromodulators^86, 87^, it is likely that Ciona’s proto-hypothalamic structure shares not only molecular but also functional similarities with its vertebrate counterpart.

### Summary and Outlook

This study shows that Ciona locomotor behavior is remarkably complex and flexibly structured, especially when we consider that the larval nervous system is equipped with less than 250 neurons. This complexity in behavioral output is likely conserved across tunicate larvae as indicated by earlier findings from two different tunicate clades the Aplousobranchia^49, 62^ and Appendicularia^19^.

Significantly, we have discovered that bioamines play a major role in the observed complexity and flexibility of the locomotor repertoire by modulating postural features, behavioural modules and their transitions, during spontaneous swimming and in response to sensory stimulation. This is in line with studies across invertebrate and vertebrate species, suggesting that bioamines have an evolutionarily conserved functional role in modulating locomotor behaviors ^29, 34, 35, 36, 45, 78, 88, 89, 90^. Furthermore, similarly to what has recently been shown in mice, our findings indicate that these behavioral modules represent a class of druggable targets^42^, facilitating future investigations.

While modern neuroscience has strongly benefited from the classic model systems, recent technological developments have encouraged the expansion of functional studies to non-traditional models^18, 91^.

We have now established a framework for a higher-throughput yet higher-resolution dissection of the behavioral repertoire of *Ciona*. Our experiments reveal that the analytical approaches we have taken are capable of systematically capturing known and new behaviors that were unidentified previously. The high sensitivity of our approach can be leveraged for extracting subtle phenotypes and mapping the contribution of individual neurons and molecules to behavioral structure through chemogenetics and genome editing. Ultimately, Ciona may serve as a key organism to identify evolutionary constraints and flexibility at multiple levels of behavioral organization and reveal fundamental principles of how molecules, neurons and circuits generate the chordate behavioral repertoire.

## Methods

### Animal Collection and rearing conditions

Gravid adult Ciona intestinalis were collected from the following site in Bergen: Døsjevika, Bildøy Marina AS, postcode 5353, Norway. The GPS coordinates of the site are as follows: 60.344330, 5.110812. Animals were kept in a purpose-built facility at 10°C with a pH of 8.2 under constant illumination. Fertilization and embryonic development conditions were as previously described^51^. Age distribution of the larvae we assayed in terms of hours post hatching is indicated in Fig. S11.

#### Drug concentrations Table

| Drug | Concentration [μM] |
| --- | --- |
| Chlorpromazine | 0.1 |
| Raclopride | 0.025 |
| Methiothepin | 1 |
| Mianserin | 0.05 |
| Fluoxetine | 1 |
| Compiramine | 10 |
| Imipramine | 10 |
| Phentolamine | 50 |
| Quinpirole | 0.03 |
| Serotonin | 100 |
| α-methyl serotonin | 5 |
| Paroxetine | 1 |
| Dopamine | 0.1 |
| Ascorbic acid | 28 |

#### Experimental setup

Each Ciona Tracker 2.0 is built using a DMK 33UP1300 (Imaging Source) coupled to an MVL75M1 lens and two C-mount extensions CML10 and CML25 (Thorlabs). Individual Ciona larvae were filmed using an array of 6 modular Ciona Tracker 2.0 systems. Each animal was assayed over a period of 30 minutes and this period was split into four recordings. An initial 15 minutes acclimatization period movie and a subsequent 3 × 5 minutes acclimatized period movies.

##### Light experiments

Light stimuli (white, Red, Green or Blue) were given for 1 minute starting at the 30^th^ second and ending at the 90^th^ second of the first 5 minutes movie. The following table details the properties of the LEDs used to delivery the stimuli

| Emitting Color | Model | Wavelength (nm) | Luminous intensity (med) | Voltage (V) |
| --- | --- | --- | --- | --- |
| Red | 13CBAUP | 620-630 | 550-700 | 1.8-2.2 |
| Green | 13CGAUP | 515-530 | 1100-1400 | 3.0-3.2 |
| Blue | 10R1MUX | 465-475 | 200-400 | 3.0-3.4 |

White light stimuli were delivered using a HALOSTAR 10 W 12 V G4 halogen lamp with a nominal luminous flux of 130 lm. Its spectral power distribution can be found here: https://docs.rs-online.com/7d94/0900766b8128288b.pdf

#### Tierpsy analysis

Videos of larvae recorded using our behavioral setup was then analyzed with the help of Tierpsy software package to extract positional data. The software segments the larval pixels from the background of the arena and identifies the two contours of the larvae. The software then calculates 49 equally spaced coordinates on the two contours such that the first pair of coordinates represents the tip of the head and the 49th pair represents the tip of the larval tail. The software also calculates the width of the larvae as the distance between the corresponding points on the two contours and uses this to calculate the midline (henceforth referred to as a skeleton) described by 49 coordinates.

#### Feature extraction

Following the Tierpsy analysis, we calculate a set of biophysical features with an aim to quantitatively describe the movement of the larvae in the arena. In order to quantify the amount by which parts of the larval body deviates from a straight line while swimming, we calculated the curvature at each of the 49 points on the skeleton. Assuming the skeleton to be a differentiable curve, the curvature was measured as the rate of change of the curve’s tangent angle with respect to its arc length(as defined in^37^). To calculate the numerical derivatives, we used a Savitzky-Golay filter of window length 15 and polynomial order 2 using the implementation in the scipy, so that the skeleton is approximated by a smooth curve.

To visualize the correlation of curvature values along the length of the larval body, we calculated the covariance matrix. Curvature from a subset of 231 experiments or animals/ larvae (where Tierpsy software successfully identified the larvae from all of the frames in the videos) with a total of 2290901 frames were used to obtain the 49 × 49 matrix (Fig1). The smooth structure of the correlation matrix was indicative of a strong correlation and hence an existence of a lower dimensional feature space. We performed an eigen decomposition of the covariance matrix (principal component analysis) to obtain the eigenvectors (principal components) and eigenvalues (explained of each of the components). The eigenvectors were sorted by the eigenvalues and the 6 top eigenvectors which explained 97% of the variance were selected to provide a lower dimensional description of the curvature of the larvae. The six eigenvectors are referred to as eigencionas in the paper. Having defined the six eigenvectors as features, we can calculate eigen coefficients EC1, EC2,.., EC6 (principal components scores) at each time point (or for each frame) which describes the posture of the skeleton.

We also calculate quirkiness as a scalar valued feature in the range of 0 to 1 indicating the eccentricity of the larval body, as explained in Tierpsy^37^. Speed at each of the 49 points of the skeleton is also calculated across time as the distance by which the skeletal point moves in the arena between two adjacent frames in the video.

We defined seven distinct body parts or segments on the larvae by grouping the 49 skeleton points. Initially, we identified a point in the range of 4 to 22 along the skeleton where the contour width decreases sharply (local minima of the derivative of contour width along the skeleton) and defined it as the neck point. The change in contour width is characteristic of the neck, where the wide head region ends and the narrower tail of the larvae starts. The neck segment is defined such that it comprises three skeleton points with the neck point as the centre. The points on the skeleton which lie anterior to this segment are hence grouped into a head segment and the coordinates that lie after are grouped into a tail segment. The skeleton points tail segment is further divided into 5 segments, namely tail_base (TB), tail_pre_mid (TprM), tail_mid (TM), tail_post_mid (TpoM), tail_tip (TT). such that a summary of movement of the tail can be obtained without limiting the degrees of freedom.

Following the definition of the 7 body segments, we calculated another postural feature, namely relative tangent angles. Initially, tangent angles were defined for each of the segments as the angle made by the line segments joining the end points of the segments (on the skeleton) with the x-axis of the video frame. We used the arctan2 function in the numpy package to compute the tangent angles (in radians) from the xy coordinates of the two end points of each of the segments. To obtain a measurement in the larvae’s coordinate system rather than the global coordinate system of the arena, we computed the difference of these tangent angles with respect to the tangent angle of the head segment. These differences were then defined as the six relative tangent angles describing the posture of the skeleton, one for each of the segments from neck to tail_tip.

#### Statistics

For each of the parameters/features and each of the experimental conditions, we tested the data for normality using the Shapiro-wilk test with an alpha value of 0.05. Since the pvalues were less than the alpha value, the null hypothesis that data is from a normal distribution was rejected. Hence for comparison between different groups in the further analysis, we used non-parametric tests. In the case of comparison of features like curvature, speed and relative tangent angles across the 6 body segments for control/wild type dataset, we used Wilcoxon signed-rank test with an alpha value of 0.05. We used this test under the assumption that feature values across segments (along the body) are dependent due to the anatomy of the larvae. Similarly, we used Wilcoxon for comparing the 6 eigen coefficient values. The pvalues from the two-tailed alternative were used to reject the null hypothesis that the median of the differences between two distributions is zero. In addition to the two-tailed test, we also computed the pvalues for one-tailed (greater and less) tests for determining which group in a pair was significantly greater or lesser than the other.

For the comparison of features between each of the drugs against the control/wild type group, we used Mann-Whitney U tests with a Bonferroni correction. The alpha value was set at 0.05/25 = 0.002 after Bonferroni correction. While the two-tailed test is used to test the alternative hypothesis that two distributions are not equal, we used one-tailed tests (greater and less) to test if one of the distributions is stochastically greater or less than the other. For Mann-Whitney U tests, we calculated the effect size by dividing the test statistic by the product of the number of samples in each of the two groups being compared. All the statistical tests were implemented in python using the scipy package. The N, p values and test statistics values for each of the tests are provided in the supplementary tables. The calculated effect sizes for Mann-Whitney U tests have also been provided in Table S33-S35. Note that all drugs except Dopamine were compared to wild-type. Dopamine was compared to ascorbic acid, which was used as a solvent for Dopamine to stabilize it and prevent oxidation.

#### Matrix Profile Methods

Motifs were identified by calculating the matrix profile of the multidimensional time-series of 7 curvature values along the skeleton of each animal, using the mstump algorithm implementation in the stumpy python library. A rolling mean filter over 10 frames was applied to the timeseries prior to matrix profiling. Recurring motifs were defined as stretches of either 30 or 150 frames with the starting point at locations where peaks in the matrix profile are under a set threshold value of .8. This resulted in a set of 87569 motifs over 30 frames, and a set of 18776 motifs over 150 frames.

The sets of motifs were subsequently clustered into 15 groups using the TimeSeriesKMeans clustering algorithm in the tslearn python library. The cluster number was decided by adding clusters until the decrease in final model inertia started levelling out.

Clusters were annotated by generating gifs of the skeletons over the duration of the motifs for each cluster, and manually confirm if there is an enrichment for a certain behavior within a cluster.

#### HMM methods

We used a simple Gaussian Hidden Markov Model (G-HMM) to model our data. A python based open-source library hmmlearn was used to implement the model and the related algorithms. We used a feature set derived from 1743 recording across multiple experimental conditions for training the model to capture a wide range of behaviors. The recordings were chosen such that Tierpsy software had successfully segmented the larvae for at least 80% of the frames. For the training the model, we chose to use the 6 eigen features and the quirkiness features from the selected experiments. Thus, our training set consisted of a set of 1743 sequences, each one of them being a 7-dimensional time series of varying lengths (durations).

The model was trained using the ‘fit’ function of the hmmlearn library. The function essentially performs an expectation maximization (EM) algorithm to estimate the parameters of the HMM model from the time series data. The learned parameters include the state transition probabilities of the fitted model and the Gaussian distributions corresponding to each of the HMM states. Based on the learned model, the most probable state sequence for each of the 1743 time series were obtained using the Viterbi algorithm implemented as the ‘predict’ function in the hmmlearn library. This enabled us to visualize the underlying state at each time point (frame) for any given time series.

Different models were trained with the number of hidden states chosen as 6, 8, 10, 12 and 15. Also, two types of covariance matrices: ‘full’ (or unrestricted) and ‘diagonal’ were tested. On a qualitative inspection, it was observed that when the number of states were chosen as 6 or 8, the active swimming behaviors were not well distinguished. On the other hand, a choice of 12 or 15 states gave rise to learning of states which appear to occur in the dataset with a frequency of less than 1% of the frames. A 10-state model with full covariance was observed to model distinct active states without over segmenting the data and enabled us to model any dependencies between eigen coefficient features and quirkiness.

The means and variances from the Gaussian distributions of each of the 10 states of the fitted model were analysed to characterize and distinguish each of the states. Following the inference of states for each of the frames in the dataset, the percentage of occurrence of each of the 10 HMM states (behavioral states) were calculated for each of the different experimental conditions (wild type, drugs, light stimuli etc). We also recalculated the probability of state transitions for each of these experimental conditions separately.

#### Spatio-temporal mapping methods

694 control videos (wild type animals) were selected for the analysis / finding a behavioral space such that Tierpsy software identified the larvae in at least 80% of the frames. (This dataset is a subset of the experiments selected for HMM). We initially calculated Morlet wavelet transformations of the 6 eigen coefficient feature vectors of these experiments, at 30 uniformly spaced frequencies in the range of 1 to 30 Hz, giving a set of 694 distinct time series each of 180 dimensions.

To prepare the input dataset, we sampled a total of 200,000 frames from 694 wild type animals (control experiments). The sampling was done such that the probability of being sampled is proportional to a weighted sum of speeds in the neck and 5 tail segments. The speed of the neck segment was assigned a 50% weightage whereas the 5 tail segments had a 10% weightage. The sampling was used to ensure that the active swimming behavior is well represented in the input dataset. The 180 wavelet feature values corresponding to the sampled frames were then used as the input features to create a wavelet feature dataset of shape 200,000 × 180.

To obtain an interpretable visualization of the high dimensional wavelet featureset, we initially calculated a two-dimensional tSNE embedding (embedding1). TSNE embeddings and associated learning algorithms were implemented using the python package openTSNE (https://opentsne.readthedocs.io/en/latest/). Embedding1 was initialized using a PCA based initialization and cosine-based metric was used for distances. A two-step optimization process was performed (using the optimize function in openTSNE) to learn the embedding1, where the first step was run with a perplexity parameter of 500, exaggeration value of 12 (early exaggeration phase) and was followed by a second phase with a reduced exaggeration value of 1.

To facilitate effective clustering and identify stereotyped behaviors from the dataset, we crafted a 4-dimensional vector using 3 distinct embeddings obtained using different parameter combinations with tSNE implementation in the python based openTSNE library. The 4-dimensional t-SNE space was created by combining the x and y dimensions of the embedding1, x dimension of embedding2 (perplexity= 250 and exaggeration=3) and y dimension of embedding3 (perplexity= 750 and exaggeration=2). This 4D space was clustered based on the density of datapoints (into 6 regions) using the pythonic implementation of DBSCAN algorithm available in scikit-learn library. The algorithm also learned an outlier class and less than 0.001 % of the data points were clustered as outliers. To assign any new or unseen data into one of the learned clusters in the behavior space, a kNN based classifier was trained with the results of the DBSCAN algorithm. We used the KNeighborsClassifier implementation in scikit learn library with n_neighbours = 200 and distance-based weights. We then followed a similar pipeline of methods to obtain clustering results for all the frames from 1743 experiments across experimental conditions using the precomputed embeddings. The clusters were then assigned to this processed dataset so that we obtain the behavioral cluster for all the frames in our dataset.

We then randomly sampled 100 skeletons for each of the clusters, translated and rotated them such that the coordinates of the neck point and the end (49th) point lie on a vertical axis (Fig 5B). Also, we visualized the trajectories of the neck points in the arena for a randomly selected set of experiments such that each point is labelled by the cluster into which it was classified. To analyze the differences in behavioral space occupied by the larvae under different experimental conditions, we computed 2D histogram smoothed by a gaussian filter from the scatter plots of the behavioral spaces. These were plotted as 2D density graphs where the intensity is set to saturate at 0.8% of the maximum intensity (Fig5D).

## Data Availability

Skeletonized data for this paper can be found here: https://zenodo.org/record/3926785#.YZt1u7oo9yE

