## Supplementary Information for "Comprehensive analysis of behavioral dynamics in the protochordate *Ciona intestinalis*"

**Supplementary Material**

**Table S1.**

Number of animals used in our analysis

| Drug | Stimulus | #Number of animals |
|---|---|---|
| None (Control) | - | 694 |
| Clomipramine | - | 59 |
| Chlorpromazine | - | 51 |
| Methiothepin | - | 77 |
| Imipramine | - | 69 |
| Mianserin | - | 39 |
| Phentolamine | - | 38 |
| Fluoxetine | - | 32 |
| Paroxetine | - | 57 |
| Alpha-Methylserotonin | - | 55 |
| Serotonin | - | 64 |
| Ascorbic acid | - | 64 |
| Dopamine | - | 88 |
| Raclopride | - | 35 |
| Quinpirole | - | 41 |
| None (Control) | Blue light | 23 |
| None (Control) | Green light | 37 |
| None (Control) | Red light | 27 |
| None (Control) | White light | 18 |

**Table S2.**

Number of video frames per condition assayed that we used in our analysis.

| Drug | Stimulus | Stimulus interval | Video frames |
|---|---|---|---|
| Wild-type | - | - | 6603509 |
| Clomipramine | - | - | 610899 |
| Chlorpromazine | - | - | 455949 |
| Methiothepin | - | - | 737811 |
| Imipramine | - | - | 700869 |
| Mianserin | - | - | 330628 |
| Phentolamine | - | - | 439041 |
| Fluoxetine | - | - | 264167 |
| Paroxetine | - | - | 465657 |
| Alpha-Methylserotonin | - | - | 480233 |
| Serotonin | - | - | 671126 |
| Ascorbic acid | - | - | 629473 |
| Dopamine | - | - | 826246 |
| Raclopride | - | - | 382973 |
| Quinpirole | - | - | 401155 |
| Wild-type | Blue light | None | 95935 |
| | | tOFF After | 1173 |
| | | tOFF Before | 5086 |
| | | tON After | 1583 |
| | | tON Before | 6148 |
| Wild-type | Green light | None | 129400 |
| | | tOFF After | 1404 |
| | | tOFF Before | 6261 |
| | | tON After | 2472 |
| | | tON Before | 10035 |
| Wild-type | Red light | None | 112242 |
| | | tOFF After | 1497 |
| | | tOFF Before | 6122 |
| | | tON After | 1810 |
| | | tON Before | 7054 |
| Wild-type | White light | None | 91238 |
| | | tOFF After | 1254 |
| | | tOFF Before | 4812 |
| | | tON After | 1191 |
| | | tON Before | 4946 |

**Table S3.**

P-values for data corresponding to Fig.1G

| Greater than | | | | | | |
|---|---|---|---|---|---|---|
| | **cN** | **cTB** | **cTprM** | **cTM** | **cTpoM** | **cTT** |
| **cN** | | 1 | 1 | 1 | 1.9E-151 | 0 |
| **cTB** | 0 | | 1.65E-57 | 0 | 0 | 0 |
| **cTprM** | 0 | 1 | | 0 | 0 | 0 |
| **cTM** | 0 | 1 | 1 | | 0 | 0 |
| **cTpoM** | 1 | 1 | 1 | 1 | | 0 |
| **cTT** | 1 | 1 | 1 | 1 | 1 | |
| Less than | | | | | | |
| | **cN** | **cTB** | **cTprM** | **cTM** | **cTpoM** | **cTT** |
| **cN** | | 0 | 0 | 0 | 1 | 1 |
| **cTB** | 1 | | 1 | 1 | 1 | 1 |
| **cTprM** | 1 | 1.65E-57 | | 1 | 1 | 1 |
| **cTM** | 1 | 0 | 0 | | 1 | 1 |
| **cTpoM** | 1.9E-151 | 0 | 0 | 0 | | 1 |
| Two-sided | | | | | | |
| | **cN** | **cTB** | **cTprM** | **cTM** | **cTpoM** | **cTT** |
| **cN** | | 0 | 0 | 0 | 3.8E-151 | 0 |
| **cTB** | 0 | | 3.3E-57 | 0 | 0 | 0 |
| **cTprM** | 0 | 3.3E-57 | | 0 | 0 | 0 |
| **cTM** | 0 | 0 | 0 | | 0 | 0 |
| **cTpoM** | 3.8E-151 | 0 | 0 | 0 | | 0 |

**Table S4.**

P-values for data corresponding to Fig.1H

| Greater than | | | | | | |
|---|---|---|---|---|---|---|
| | **rtaN** | **rtaTB** | **rtaTprM** | **rtaTM** | **rtaTpoM** | **rtaTT** |
| **rtaN** | | 1 | 1 | 1 | 1 | 1 |
| **rtaTB** | 0 | | 1 | 1 | 1 | 1 |
| **rtaTprM** | 0 | 0 | | 1 | 1 | 1 |
| **rtaTM** | 0 | 0 | 0 | | 1 | 1 |
| **rtaTpoM** | 0 | 0 | 0 | 0 | | 0 |
| **rtaTT** | 0 | 2.3E-102 | 0 | 0 | 1 | |
| Less than | | | | | | |
| | **rtaN** | **rtaTB** | **rtaTprM** | **rtaTM** | **rtaTpoM** | **rtaTT** |
| **rtaN** | | 0 | 0 | 0 | 0 | 0 |
| **rtaTB** | 1 | | 0 | 0 | 0 | 2.3E-102 |
| **rtaTprM** | 1 | 1 | | 0 | 0 | 0 |
| **rtaTM** | 1 | 1 | 1 | | 0 | 0 |
| **rtaTpoM** | 1 | 1 | 1 | 1 | | 1 |
| Two-sided | | | | | | |
| | **rtaN** | **rtaTB** | **rtaTprM** | **rtaTM** | **rtaTpoM** | **rtaTT** |
| **rtaN** | | 0 | 0 | 0 | 0 | 0 |
| **rtaTB** | 0 | | 0 | 0 | 0 | 4.6E-102 |
| **rtaTprM** | 0 | 0 | | 0 | 0 | 0 |
| **rtaTM** | 0 | 0 | 0 | | 0 | 0 |
| **rtaTpoM** | 0 | 0 | 0 | 0 | | 0 |

**Table S5.**

P-values for data corresponding to Fig.1I

| Greater than | | | | | | |
|---|---|---|---|---|---|---|
| | **sN** | **sTB** | **sTprM** | **sTM** | **sTpoM** | **sTT** |
| **sN** | | 1 | 1 | 1 | 0 | 0 |
| **sTB** | 0 | | 1 | 1 | 0 | 0 |
| **sTprM** | 0 | 0 | | 0 | 0 | 0 |
| **sTM** | 0 | 0 | 1 | | 0 | 0 |
| **sTpoM** | 1 | 1 | 1 | 1 | | 0 |
| **sTT** | 1 | 1 | 1 | 1 | 1 | |
| Less than | | | | | | |
| | **sN** | **sTB** | **sTprM** | **sTM** | **sTpoM** | **sTT** |
| **sN** | | 0 | 0 | 0 | 1 | 1 |
| **sTB** | 1 | | 0 | 0 | 1 | 1 |
| **sTprM** | 1 | 1 | | 1 | 1 | 1 |
| **sTM** | 1 | 1 | 0 | | 1 | 1 |
| **sTpoM** | 0 | 0 | 0 | 0 | | 1 |
| Two-sided | | | | | | |
| | **sN** | **sTB** | **sTprM** | **sTM** | **sTpoM** | **sTT** |
| **sN** | | 0 | 0 | 0 | 0 | 0 |
| **sTB** | 0 | | 0 | 0 | 0 | 0 |
| **sTprM** | 0 | 0 | | 0 | 0 | 0 |
| **sTM** | 0 | 0 | 0 | | 0 | 0 |
| **sTpoM** | 0 | 0 | 0 | 0 | | 0 |

**Table S6.**

P-values for data corresponding to Fig.1P

| Greater than | | | | | | |
|---|---|---|---|---|---|---|
| | **EC1** | **EC2** | **EC3** | **EC4** | **EC5** | **EC6** |
| **EC1** | | 0 | 1 | 0 | 0 | 0 |
| **EC2** | 1 | | 1 | 1 | 1 | 1 |
| **EC3** | 0 | 0 | | 0 | 0 | 0 |
| **EC4** | 1 | 0 | 1 | | 0 | 0 |
| **EC5** | 1 | 0 | 1 | 1 | | 1 |
| **EC6** | 1 | 0 | 1 | 1 | 0 | |
| Less than | | | | | | |
| | **EC1** | **EC2** | **EC3** | **EC4** | **EC5** | **EC6** |
| **EC1** | | 1 | 0 | 1 | 1 | 1 |
| **EC2** | 0 | | 0 | 0 | 0 | 0 |
| **EC3** | 1 | 1 | | 1 | 1 | 1 |
| **EC4** | 0 | 1 | 0 | | 1 | 1 |
| **EC5** | 0 | 1 | 0 | 0 | | 0 |
| Two-sided | | | | | | |
| | **EC1** | **EC2** | **EC3** | **EC4** | **EC5** | **EC6** |
| **EC1** | | 0 | 0 | 0 | 0 | 0 |
| **EC2** | 0 | | 0 | 0 | 0 | 0 |
| **EC3** | 0 | 0 | | 0 | 0 | 0 |
| **EC4** | 0 | 0 | 0 | | 0 | 0 |
| **EC5** | 0 | 0 | 0 | 0 | | 0 |

**Movie S1.**
Video of 1-second-long sequences of skeletons from cluster A1.

**Movie S2.**
Video of 1-second-long sequences of skeletons from cluster B1.

**Movie S3.**
Video of 1-second-long sequences of skeletons from cluster C1.

**Movie S4.**
Video of 1-second-long sequences of skeletons from cluster D1.

**Movie S5.**
Video of 1-second-long sequences of skeletons from cluster E1.

**Movie S6.**
Video of 1-second-long sequences of skeletons from cluster F1.

**Movie S7.**
Video of 1-second-long sequences of skeletons from cluster G1.

**Movie S8.**
Video of 1-second-long sequences of skeletons from cluster H1.

**Movie S9.**
Video of 1-second-long sequences of skeletons from cluster I1.

**Movie S10.**
Video of 1-second-long sequences of skeletons from cluster J1.

**Movie S11.**
Video of 1-second-long sequences of skeletons from cluster K1.

**Movie S12.**
Video of 1-second-long sequences of skeletons from cluster L1.

**Movie S13.**
Video of 1-second-long sequences of skeletons from cluster M1.

**Movie S14.**
Video of 1-second-long sequences of skeletons from cluster N1.

**Movie S15.**
Video of 1-second-long sequences of skeletons from cluster O1.

**Movie S16.**
Video of 5-second-long sequences of skeletons from cluster a5.

**Movie S17.**
Video of 5-second-long sequences of skeletons from cluster b5.

**Movie S18.**
Video of 5-second-long sequences of skeletons from cluster c5.

**Movie S19.**
Video of 5-second-long sequences of skeletons from cluster d5.

**Movie S20.**
Video of 5-second-long sequences of skeletons from cluster e5.

**Movie S21.**
Video of 5-second-long sequences of skeletons from cluster f5.

**Movie S22.**
Video of 5-second-long sequences of skeletons from cluster g5.

**Movie S23.**
Video of 5-second-long sequences of skeletons from cluster h5.

**Movie S24.**
Video of 5-second-long sequences of skeletons from cluster i5.

**Movie S25.**
Video of 5-second-long sequences of skeletons from cluster j5.

**Movie S26.**
Video of 5-second-long sequences of skeletons from cluster k5.

**Movie S27.**
Video of 5-second-long sequences of skeletons from cluster l5.

**Movie S28.**
Video of 5-second-long sequences of skeletons from cluster m5.

**Movie S29.**
Video of 5-second-long sequences of skeletons from cluster n5.

**Movie S30.**
Video of 5-second-long sequences of skeletons from cluster o5.

**Movie S31.**
Animations of skeleton movements in the arena for HMM Cluster α.

**Movie S32.**
Animations of skeleton movements in the arena for HMM Cluster β.

**Movie S33.**
Animations of skeleton movements in the arena for HMM Cluster γ.

**Movie S34.**
Animations of skeleton movements in the arena for HMM Cluster δ.

**Movie S35.**
Animations of skeleton movements in the arena for HMM Cluster ε.

**Movie S36.**
Animations of skeleton movements in the arena for HMM Cluster ζ.

**Movie S37.**
Animations of skeleton movements in the arena for HMM Cluster η.

**Movie S38.**
Animations of skeleton movements in the arena for HMM Cluster θ.

**Movie S39.**
Animations of skeleton movements in the arena for HMM Cluster ι.

**Movie S40.**
Animations of skeleton movements in the arena for HMM Cluster κ.

**Movie S41.**
Animations of skeleton movements in the arena for tSNE Cluster 1.

**Movie S42.**
Animations of skeleton movements in the arena for tSNE Cluster 2.

**Movie S43.**
Animations of skeleton movements in the arena for tSNE Cluster 3.

**Movie S44.**
Animations of skeleton movements in the arena for tSNE Cluster 4.

**Movie S45.**
Animations of skeleton movements in the arena for tSNE Cluster 5.

**Movie S46.**
Animations of skeleton movements in the arena for tSNE Cluster 6.

**Movie S47.**
Animation of an animal exploring the tSNE behavioral space.

**Table S7. (separate file)**
Fig. 2 SMD values

**Table S8. (separate file)**
Mean values of violin plots for different drugs and features shown in Fig. S1 and S2.

**Table S9. (separate file)**
Standard deviation values of violin plots for different drugs and features shown in Fig. S1 and S2.

**Table S10. (separate file)**
Shapiro-Wilk tests for normality analysis of the data shown in Fig. S1 and S2.

**Table S11. (separate file)**
Mann-Whitney U two-sided tests with Bonferroni correction for statistical significance analysis of the data shown in Fig. S1 and S2.

**Table S12. (separate file)**
Mann-Whitney U one-sided (less) tests with Bonferroni correction for statistical significance analysis of the data shown in Fig. S1 and S2.

**Table S13. (separate file)**
Mann-Whitney U one-sided (greater) tests with Bonferroni correction for statistical significance analysis of the data shown in Fig. S1 and S2.

**Table S14. (separate file)**
Quantification of 1 second time-window motif clusters representation (shown as %) in wild-type and drug datasets corresponding to heat maps shown in Fig. 3F and Fig. S3E (in this case Raclopride is not included in the heat map).

**Table S15. (separate file)**

Quantification of percentage fold increase of 1 second time-window motif clusters representation of drugs relative to wild-type corresponding to heat map shown in Fig. 3G and Fig. S3G (Raclopride is omitted in this heat map). Dopamine is compared to Ascorbic acid in which it was dissolved.

**Table S16. (separate file)**

Quantification of 5 seconds time-window motif clusters representation (shown as %) in wild-type and drug datasets corresponding to heat map shown in Fig. 3H and Fig. S3F (with Raclopride omitted in this heat map).

**Table S17. (separate file)**

Quantification of percentage fold increase of 5 seconds time-window motif clusters representation of drugs relative to wild-type corresponding to the heat map shown in Fig. 3I and Fig. S3H (where Raclopride is left out). Dopamine is compared to Ascorbic acid in which it was dissolved.

**Table S18. (separate file)**

Percentage of representation of the HMM states for different drug datasets, corresponding to panels Fig. 4 E and Fig. S5 A.

**Table S19. (separate file)**

Percentage fold change of representation of the HMM states for different drug datasets, corresponding to panels Fig. 4 E and Fig. S5 B.

**Table S20. (separate file)**

Percentage of representation of the tSNE clusters for different drug datasets, corresponding to heatmap Fig. 5 G.

**Table S21. (separate file)**

Percentage fold change of representation of the tSNE clusters for different drug datasets, corresponding to panel Fig. 5H

**Table S22. (separate file)**

SMD values corresponding to Fig. 6B.

**Table S23. (separate file)**

SMD values corresponding to Fig. 6C.

**Table S24. (separate file)**

Mean values of violin plots for different color light ON stimuli shown in Fig.S8.

**Table S25. (separate file)**

Standard deviation values of violin plots for different color light ON stimuli shown in Fig.S8

**Table S26. (separate file)**
Mean values of violin plots for different color light OFF stimuli shown in Fig.S9.

**Table S27. (separate file)**
Standard deviation values of violin plots for different color light OFF stimuli shown in Fig.S9.

**Table S28. (separate file)**
Shapiro-Wilk tests for normality analysis of the data shown in Fig. S8 and S9.

**Table S29. (separate file)**
Mann-Whitney U two-sided tests with Bonferroni correction for statistical significance analysis of the data shown in Fig. S8 and S9.

**Table S30. (separate file)**
Mann-Whitney U one-sided (less) tests with Bonferroni correction for statistical significance analysis of the data shown in Fig. S8 and S9.

**Table S31. (separate file)**
Mann-Whitney U one-sided (greater) tests with Bonferroni correction for statistical significance analysis of the data shown in Fig. S8 and S9.

**Table S32. (separate file)**
Percentage of representation of the tSNE clusters for different light color stimuli ON and OFF periods, corresponding to Fig. 6 D-G, Fig. S10A-P.

**Table S33. (separate file)**
Effect size for Mann-Whitney U two-sided, corresponding to Fig. S1 and S2.

**Table S34. (separate file)**
Effect size less for Mann-Whitney U, corresponding to Fig. S1 and S2.

**Table S35. (separate file)**
Effect size greater for Mann-Whitney U, corresponding to Fig. S1 and S2.

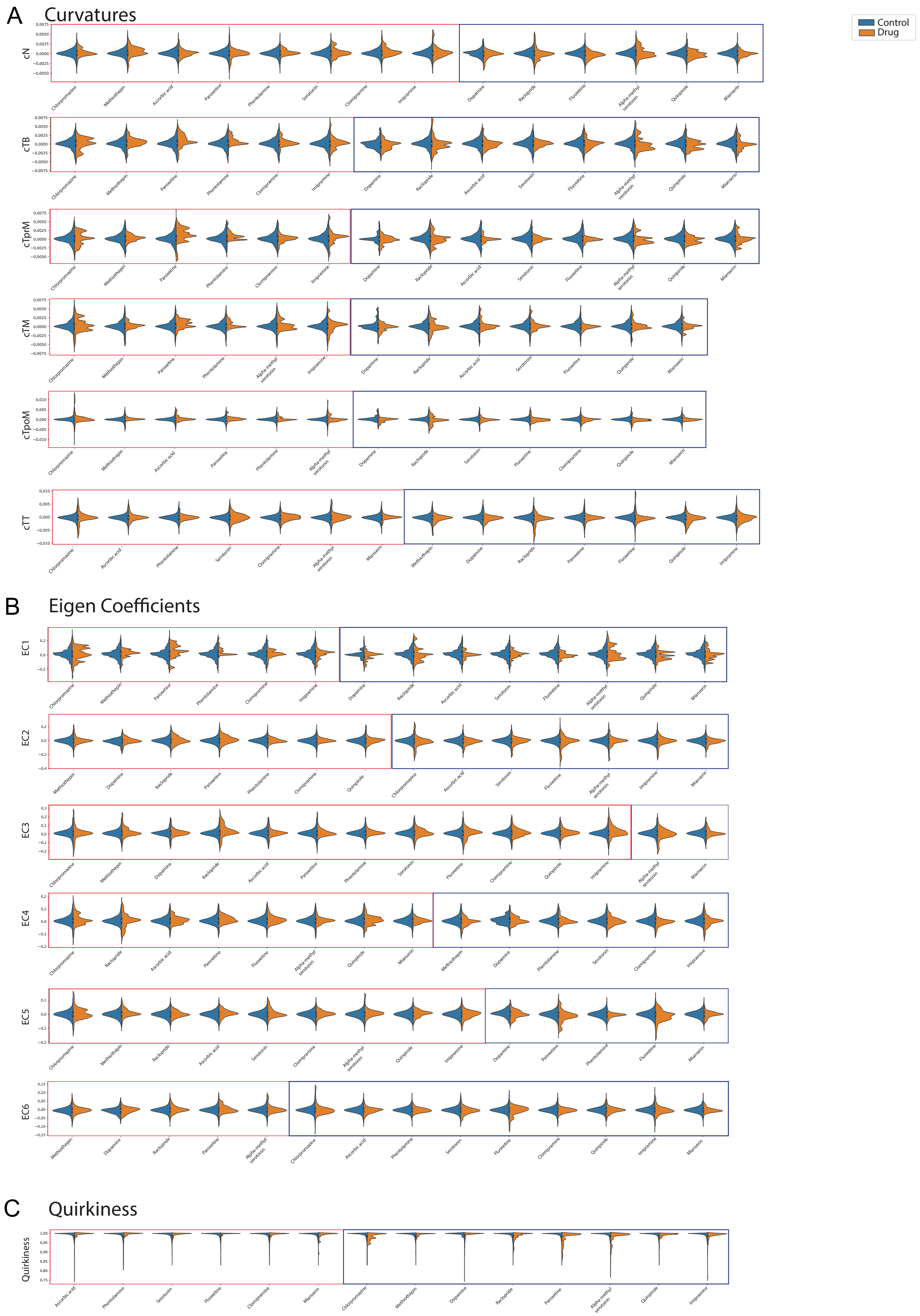

A
Curvatures
Control
Drug
cN
cTB
cTprM
cTM
cTpoM
cTT
B
Eigen Coefficients
EC1
EC2
EC3
EC4
EC5
EC6
C
Quirkiness
Quirkiness

**Fig. S1.**

(A)Violin plots comparing the distribution of curvatures of 6 body segments for different drugs with wild type. Only drugs with significant differences are shown. (B) Violin plots showing the distribution of the six eigen coefficient features for different drugs in comparison with wild type. (C) Comparison of distribution of quirkiness values exhibited in the presence of different drugs with the wild type. Note that in all the plots (A-C), drugs which showed significantly higher values of the feature are grouped together within a red-colored border whereas drugs with significantly lower feature values with respect to wild-type were grouped within a blue border. We tested for normality using the Shapiro-Wilk test ($\alpha$=0.05) (Table S10). Wild-type and drug data features were compared using Mann-Whitney U tests with a Bonferroni correction ($\alpha$=0.002) (Tables S11-S13).

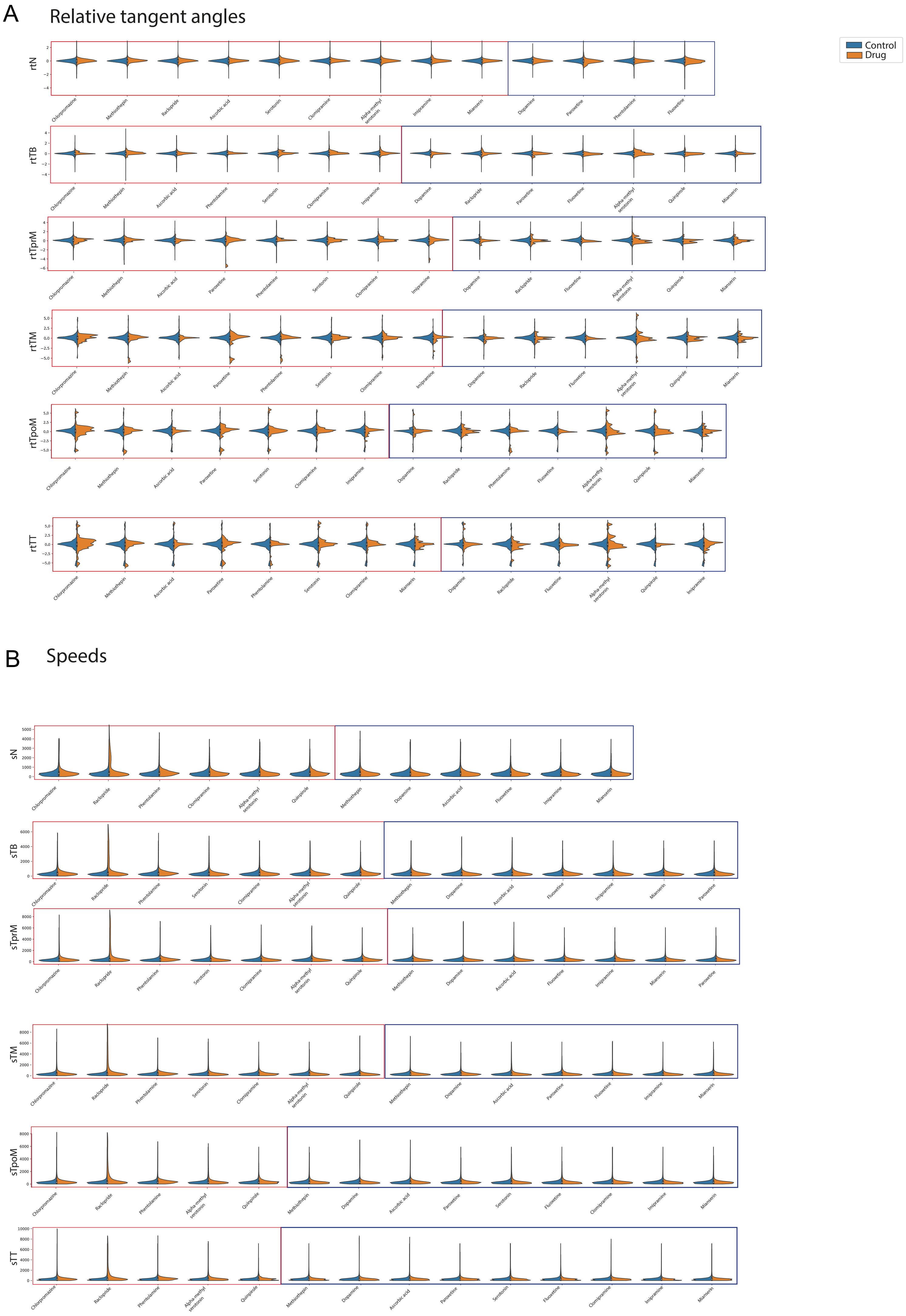
A
Relative tangent angles
Control
Drug
rtN
rtTB
rtTprM
rtTM
rtTpoM
rtTT
B
Speeds
sN
sTB
sTprM
sTM
sTpoM
sTT

**Fig. S2.**

(A)Violin plots comparing the distribution of relative tangent angles of 6 body segments for different drugs with wild type. Only drugs with significant differences are shown. (B) Comparison of distribution of speed values exhibited in the presence of different drugs with the wild type. Note that in all the plots (A,B), drugs which showed significantly higher values of the feature are grouped together within a red-colored border whereas drugs with significantly lower feature values with respect to wild-type were grouped within a blue border. We tested for normality using the Shapiro-Wilk test ($\alpha$=0.05) (Table S10). Wild-type and drug data features were compared using Mann-Whitney U tests with a Bonferroni correction ($\alpha$=0.002) (Tables S11-S13).

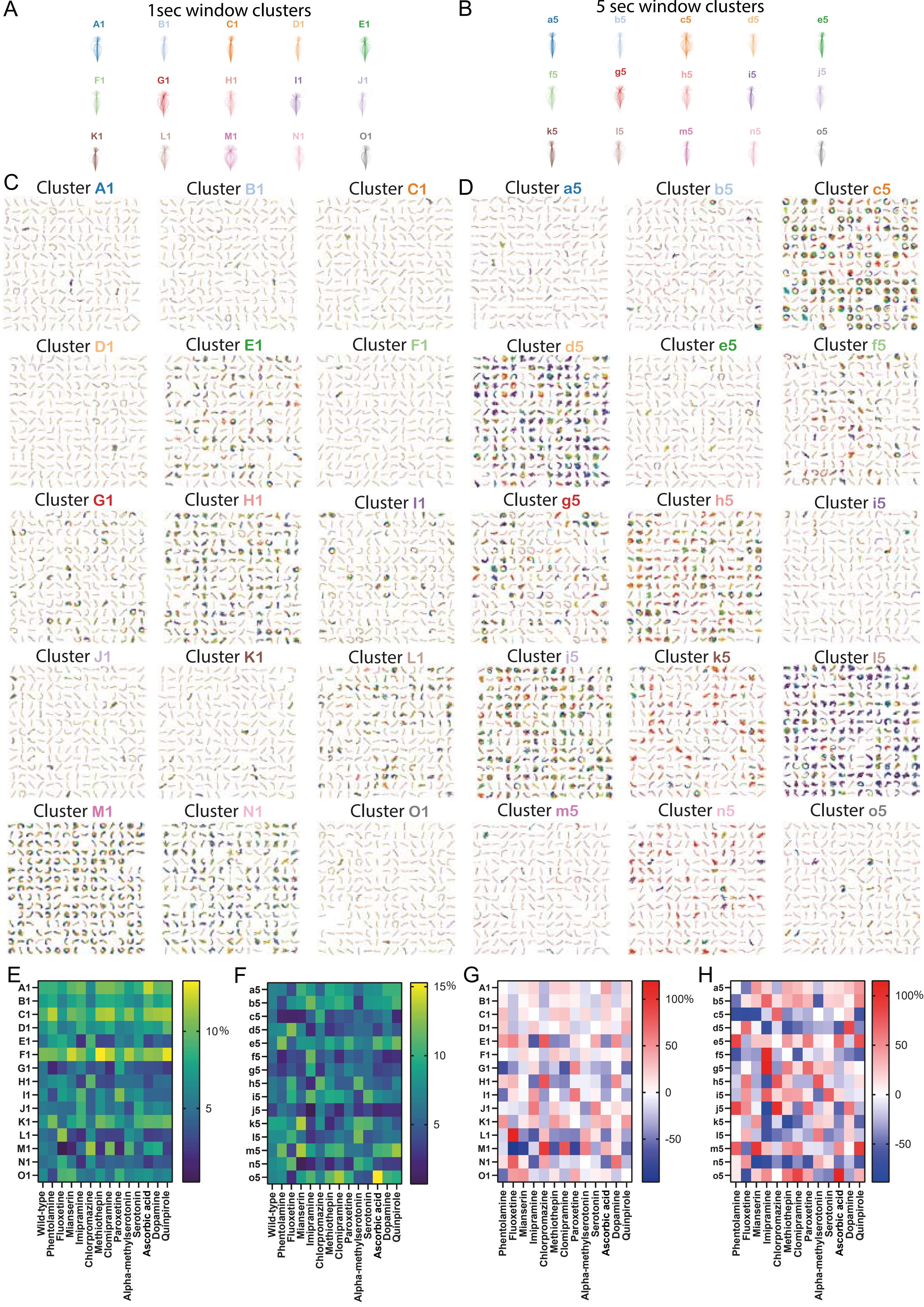

A
1sec window clusters
A1 B1 C1 D1 E1 F1 G1 H1 I1 J1 K1 L1 M1 N1 O1
B
5 sec window clusters
a5 b5 c5 d5 e5 f5 g5 h5 i5 j5 k5 l5 m5 n5 o5
C
Cluster A1
Cluster B1
Cluster C1
Cluster D1
Cluster E1
Cluster F1
Cluster G1
Cluster H1
Cluster I1
Cluster J1
Cluster K1
Cluster L1
Cluster M1
Cluster N1
Cluster O1
D
Cluster a5
Cluster b5
Cluster c5
Cluster d5
Cluster e5
Cluster f5
Cluster g5
Cluster h5
Cluster i5
Cluster j5
Cluster k5
Cluster l5
Cluster m5
Cluster n5
Cluster o5
E
10%
5
F
15%
10
5
G
100%
50
0
-50
H
100%
50
0
-50
Wild-type
Phentolamine
Fluoxetine
Mianserin
Imipramine
Chlorpromazine
Methiothepin
Clomipramine
Paroxetine
Alpha-methylserotonin
Serotonin
Ascorbic acid
Dopamine
Quinpirole

**Fig. S3.**
(A-B) Randomly selected skeletons of animals that correspond to each of the clusters shown in Fig 3 panels D and E. (C-D) Individual motifs grouped according to the motif cluster the correspond to. Panel C includes 1 second clusters, while panel D shows 5 second clusters. Motifs are color coded to show temporal progression (start→end; violet→red). (E-H) Raclopride shows much stronger effects on several clusters relative to the other drugs used in our screen. This means that in heatmaps Fig. 3F-I, it is hard to visually appreciate the differences between wild type and the other drugs. Thus, in panels E-H we show the same heatmaps but we exclude Raclopride. (E)  1second time-window motif clusters representation. (F) Percentage fold changes relative to control for 1second time-window motif clusters. (G) 5 seconds time-window motif clusters representation. (H) Percentage fold changes relative to control for 5 seconds time-window motif clusters.

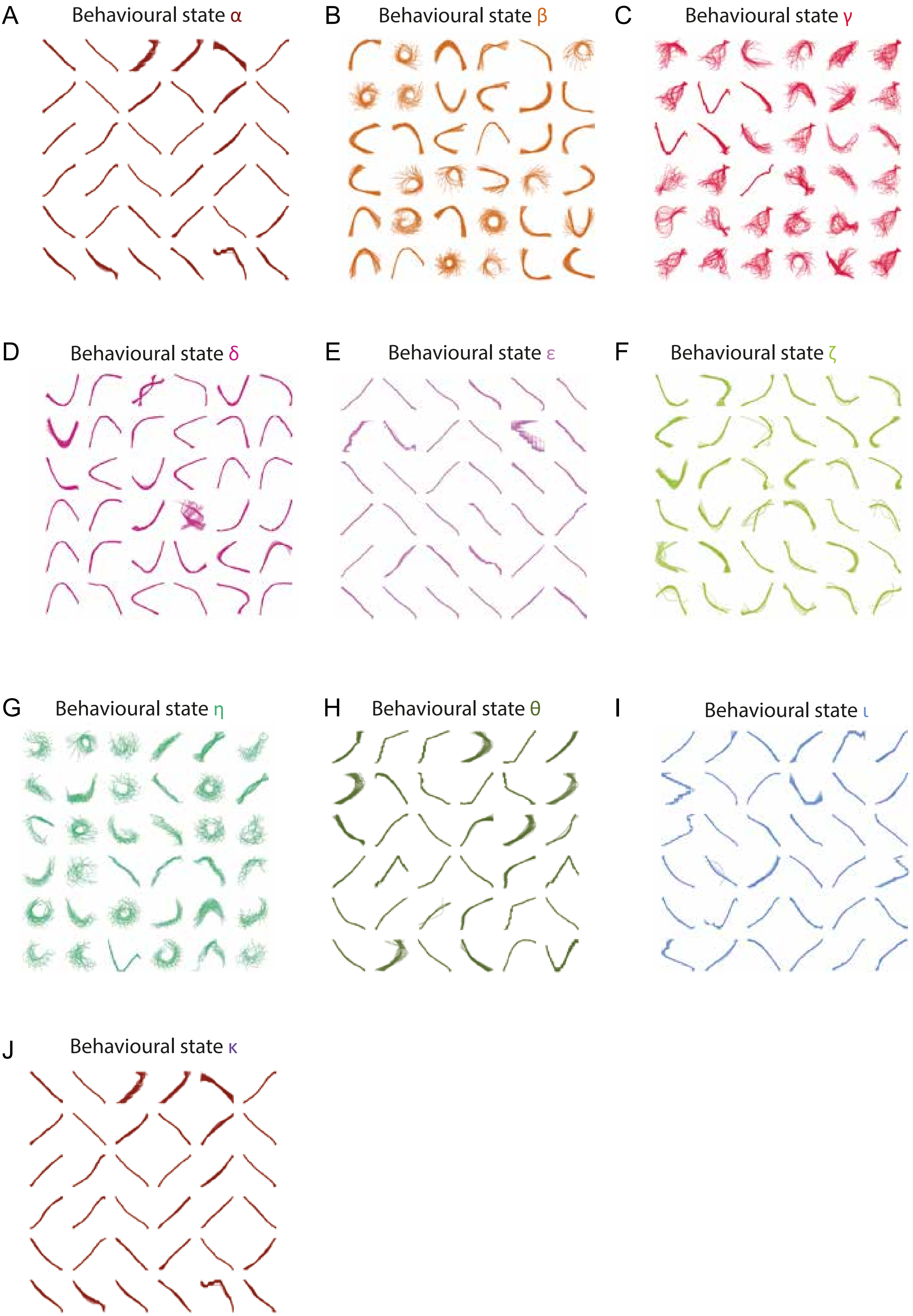
A
Behavioural state α
B
Behavioural state β
C
Behavioural state γ
D
Behavioural state δ
E
Behavioural state ε
F
Behavioural state ζ
G
Behavioural state η
H
Behavioural state θ
I
Behavioural state ι
J
Behavioural state κ

**Fig. S4.**

(A-J) Shows 36 distinct samples of skeleton trajectories exhibiting each of the 10 behavioral states (identified by HMM). Each of the 36 motifs in the panels are of a minimum of 21 frames long. These correspond to Movie S31- S40.

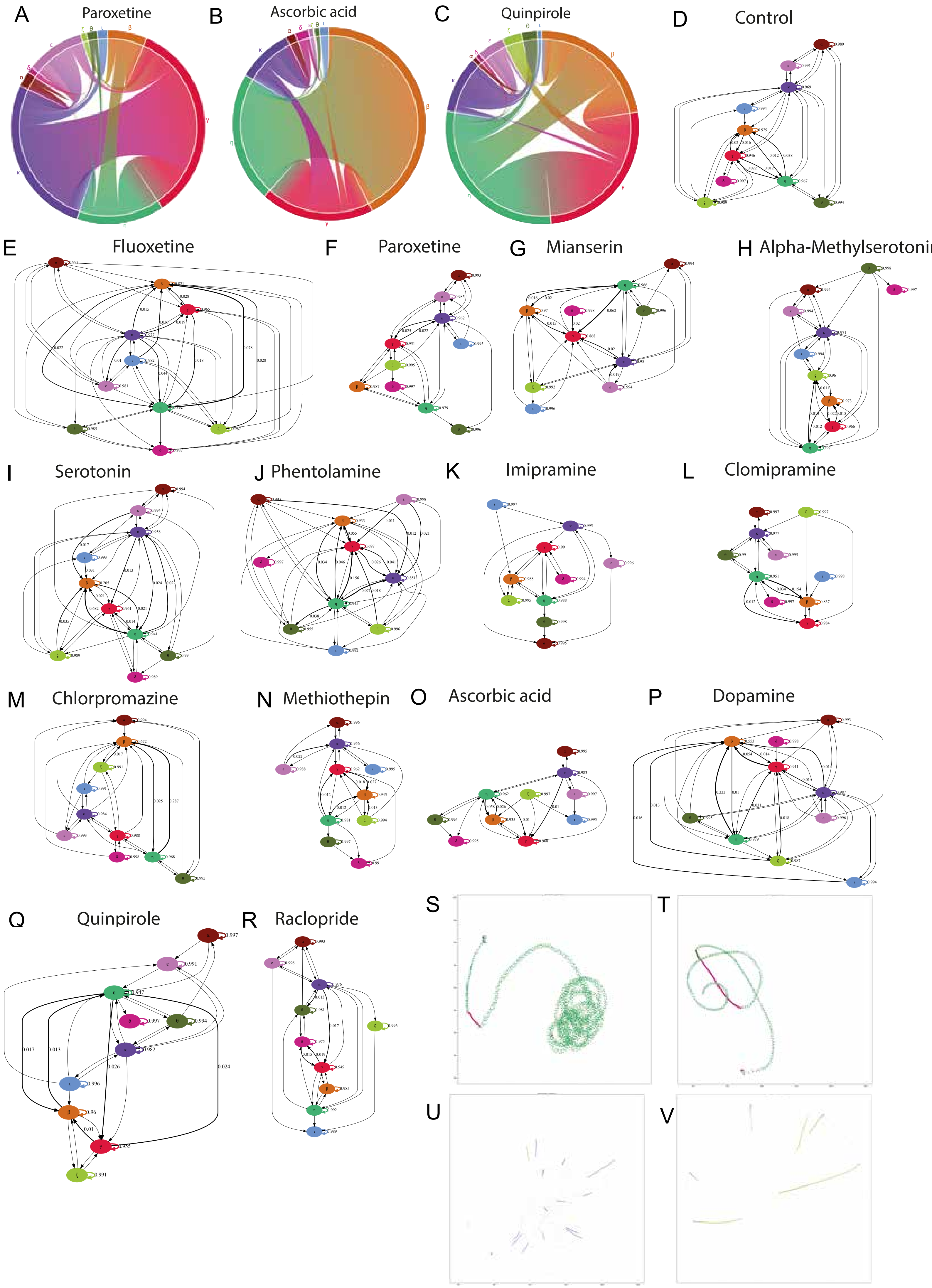
A Paroxetine
B Ascorbic acid
C Quinpirole
D Control
E Fluoxetine
F Paroxetine
G Mianserin
H Alpha-Methylserotoni
I Serotonin
J Phentolamine
K Imipramine
L Clomipramine
M Chlorpromazine
N Methiothepin
O Ascorbic acid
P Dopamine
Q Quinpirole
R Raclopride
S
T
U
V

**Fig. S5.**

(A-C) Chord diagrams showing behavioral state transitions inferred from HMM for (A) Paroxetine, (B) Ascorbic acid, (C) Quinpirole. (D-R) Markov transition graphs for wild type and drugs. The graph represents the probability of transitions from one behavioral state to any other state as defined by the HMM model. Each of the nodes in the graph represents a behavioral state. Self-transitions are represented by arrows with matching colors. Any transition with a probability greater than 0.001 are shown with an arrow. Probability values are printed for only those transitions which have a probability greater than 0.01. (S-V) Four example tracks of the neck point of larvae in the arena colored according to the behavioral state identified by HMM.

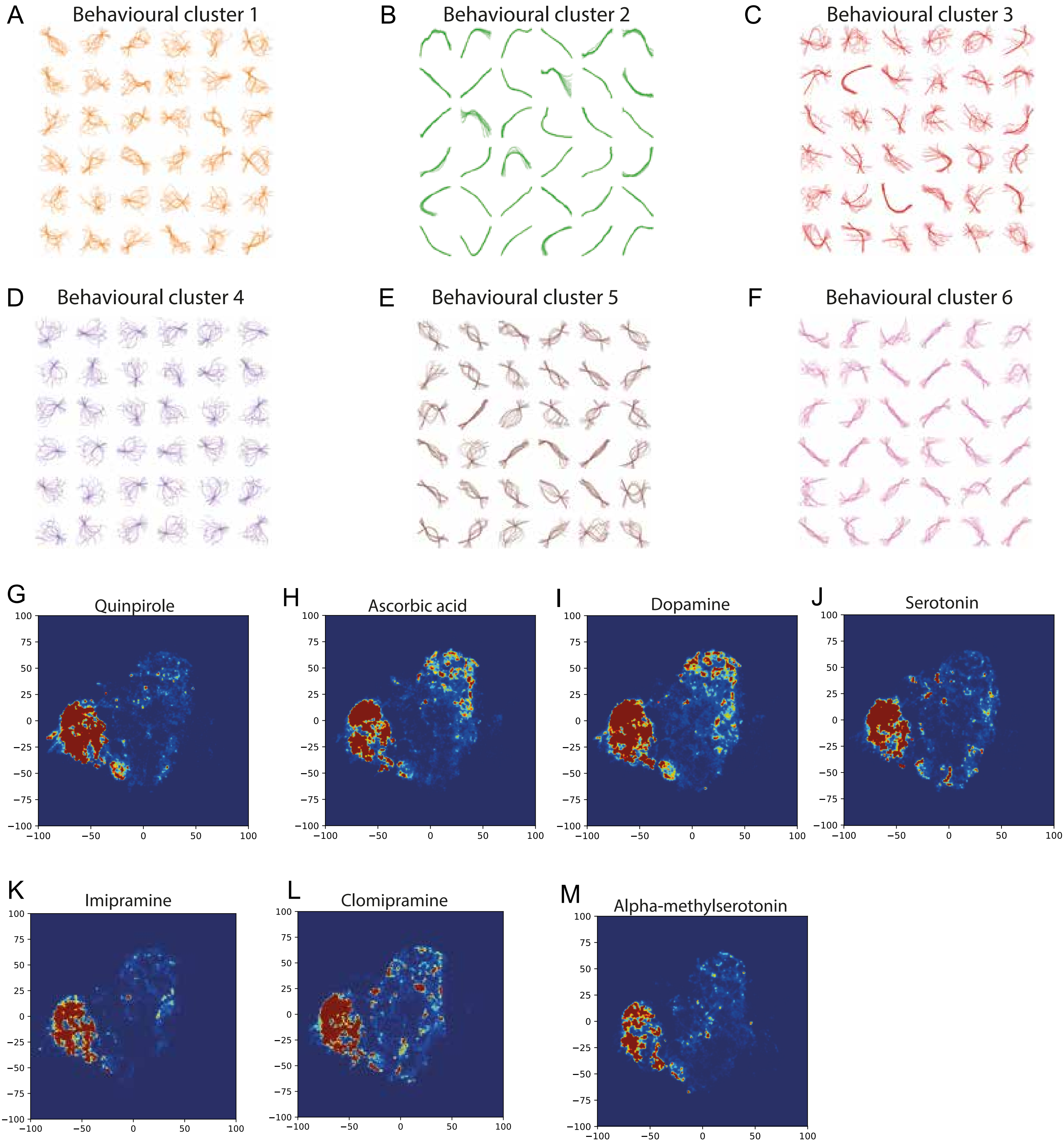

A
Behavioural cluster 1
B
Behavioural cluster 2
C
Behavioural cluster 3
D
Behavioural cluster 4
E
Behavioural cluster 5
F
Behavioural cluster 6
G
Quinpirole
H
Ascorbic acid
I
Dopamine
J
Serotonin
K
Imipramine
L
Clomipramine
M
Alpha-methylserotonin

**Fig. S6.**
(A-F) These panels show 36 distinct samples of skeleton trajectories exhibiting each of the 6 behavioral clusters inferred from the spatio-temporal embedding approach. Each of the 36 skeleton trajectories in the panels are of a minimum of 21 frames long. They correspond to Movies S41-S46. (G-M) 2D tSNE embedding of different drugs. The color is showing the density with which the different clusters are occupied (blue being lower and red higher).

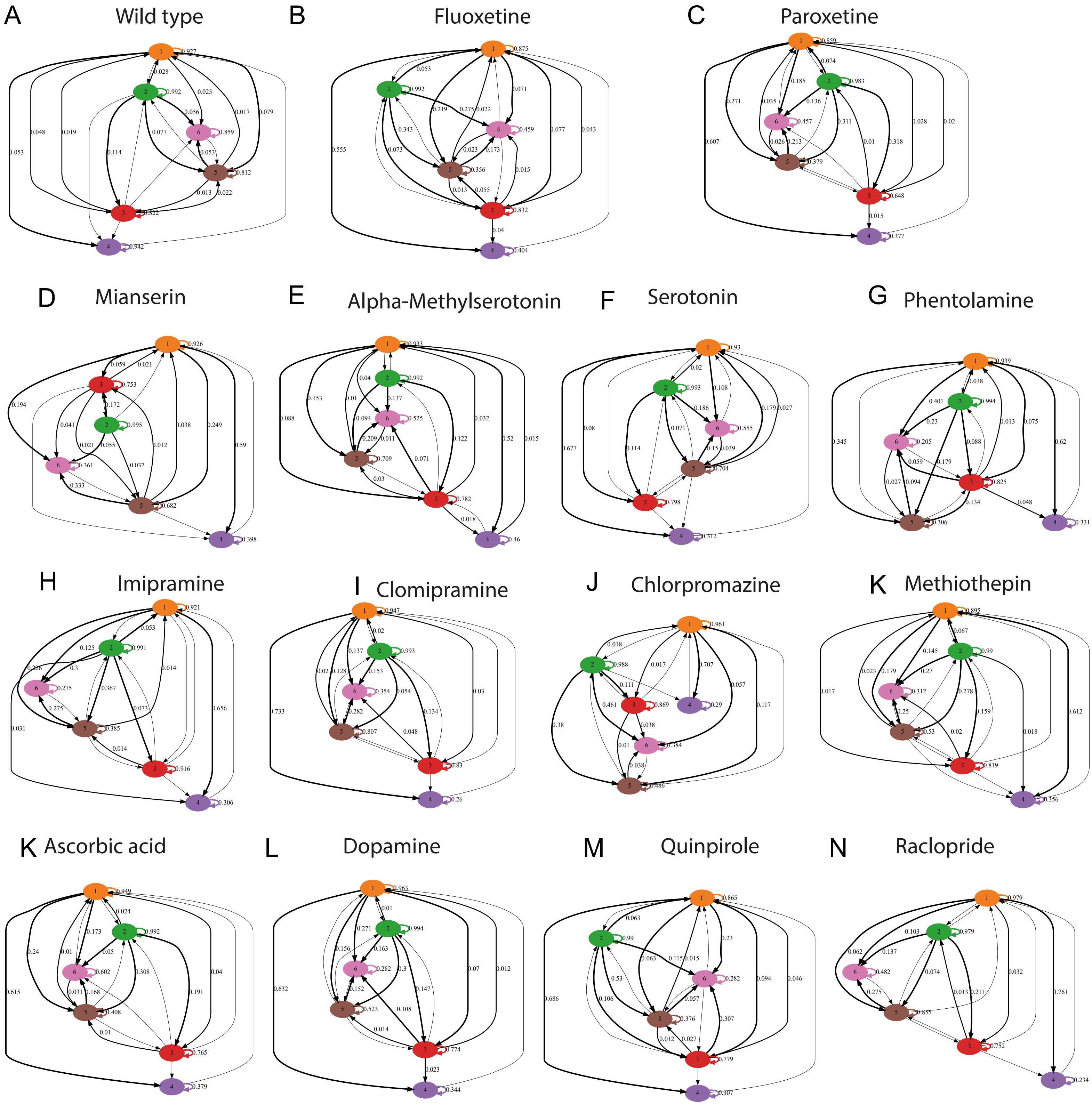
A Wild type
B Fluoxetine
C Paroxetine
D Mianserin
E Alpha-Methylserotonin
F Serotonin
G Phentolamine
H Imipramine
I Clomipramine
J Chlorpromazine
K Methiothepin
K Ascorbic acid
L Dopamine
M Quinpirole
N Raclopride

**Fig. S7.**
(A-N) Transition graphs for wild type and drugs. The graph shows the transitions from one behavioral cluster to any other cluster in terms of the probability of the transition as determined by our data. Each of the nodes in the graph represents a behavioral cluster. Self-transitions are represented by arrows with matching colors. All transitions with a probability greater than 0.001 are shown with an arrow. Probability values are printed for only those transitions which have a probability greater than 0.01.

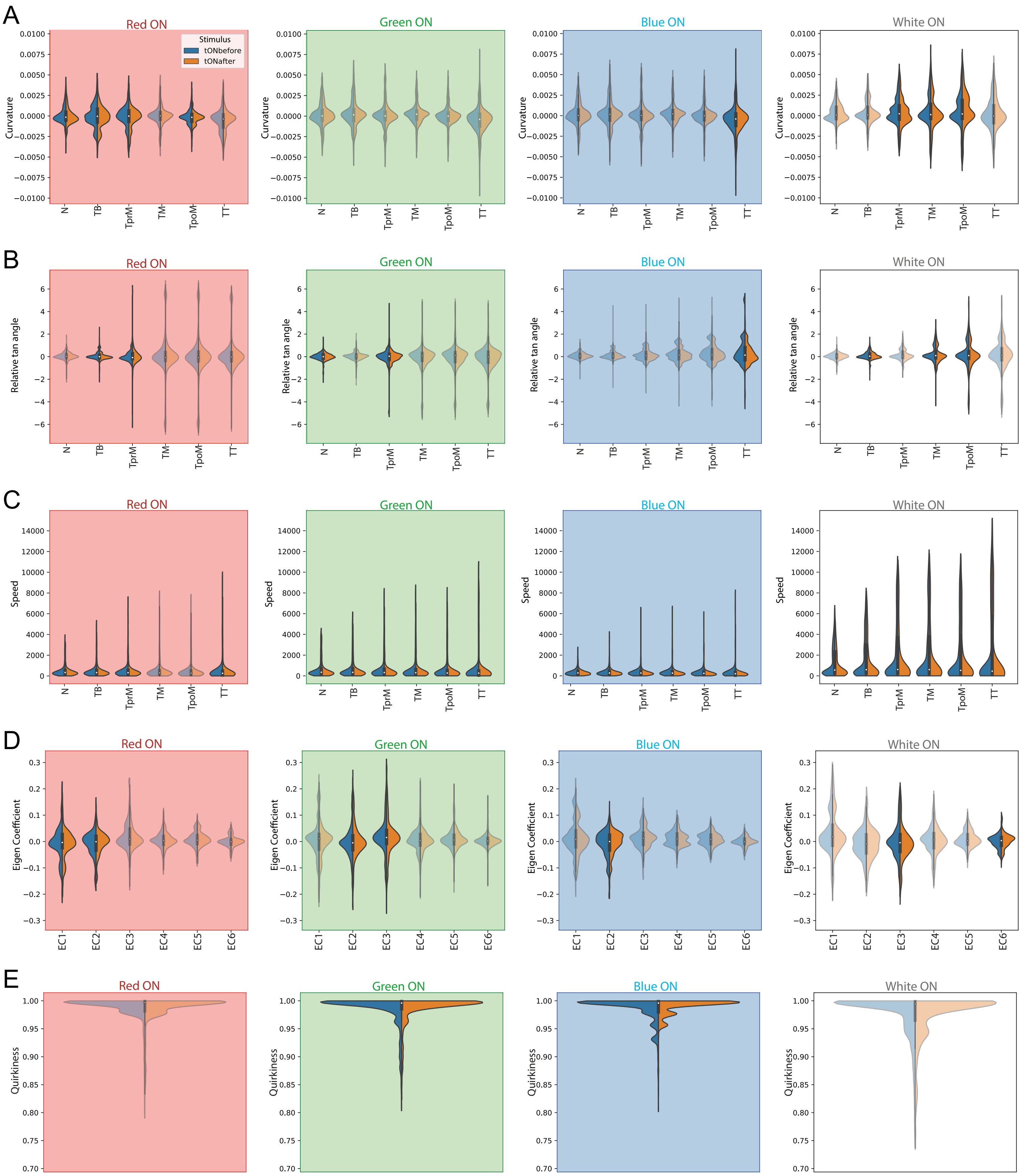
A
B
C
D
E
Red ON
Green ON
Blue ON
White ON
Stimulus
tONbefore
tONafter
Curvature
Relative tan angle
Speed
Eigen Coefficient
Quirkiness
N
TB
TprM
TM
TpoM
TT
EC1
EC2
EC3
EC4
EC5
EC6

**Fig. S8.**
(A -D) Change induced in different feature values by switching ON the light stimulus: Comparison of distribution of curvature, relative tangent angles, speed, eigen coefficient and quirkiness features before (tONbefore) and after (tONafter) the light ON event. Effect of 4 different stimuli - red, green, blue and white shown from left to right. Note: In all the panels, significant results are highlighted with a higher opacity. Tables S24-S25 provide the mean and standard deviation values for all plots shown in this figure. Tables S28-S31 provide statistical analysis for this figure. We used Shapiro-Wilk test for normality analysis. Subsequently we performed Mann-Whitney U tests with Bonferonni correction.

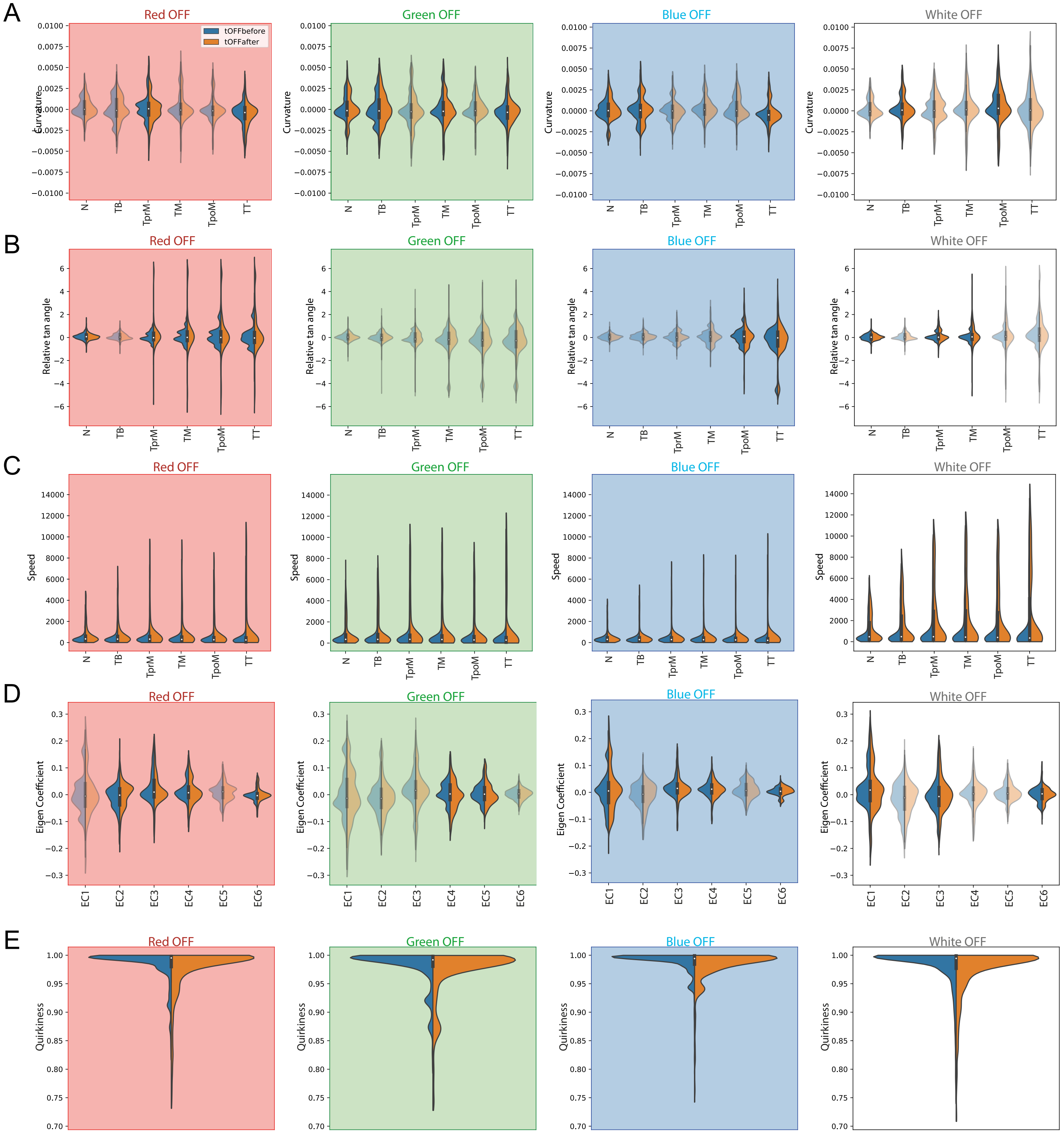
A
B
C
D
E
Red OFF
Green OFF
Blue OFF
White OFF
tOFFbefore
tOFFafter
Curvature
Relative tan angle
Speed
Eigen Coefficient
Quirkiness
N
TB
TprM
TM
TpoM
TT
EC1
EC2
EC3
EC4
EC5
EC6

**Fig. S9.**

(A) Change induced in terms of curvature values by switching OFF light stimulus: Comparison of distribution of segment-wise curvature values before (tOFFbefore) and after (tOFFafter) the light OFF event. Effect of 4 different stimuli - red, green, blue and white shown from left to right. (B-C) Plots similar to A for relative tangent angles and speeds respectively. (D) Effect of switching OFF the light stimuli in terms of change in six eigen coefficient features. (E) Change in quirkiness feature with the light OFF event. Note: In all the panels, significant results are highlighted with a higher opacity. Tables S26-S27 provide the mean and standard deviation values for all plots shown in this figure. Statistical analysis for data included in this figure can be found in Tables S28-S31. Briefly, we used Shapiro-Wilk test for normality analysis. Subsequently we performed Mann-Whitney U tests with Bonferonni correction.

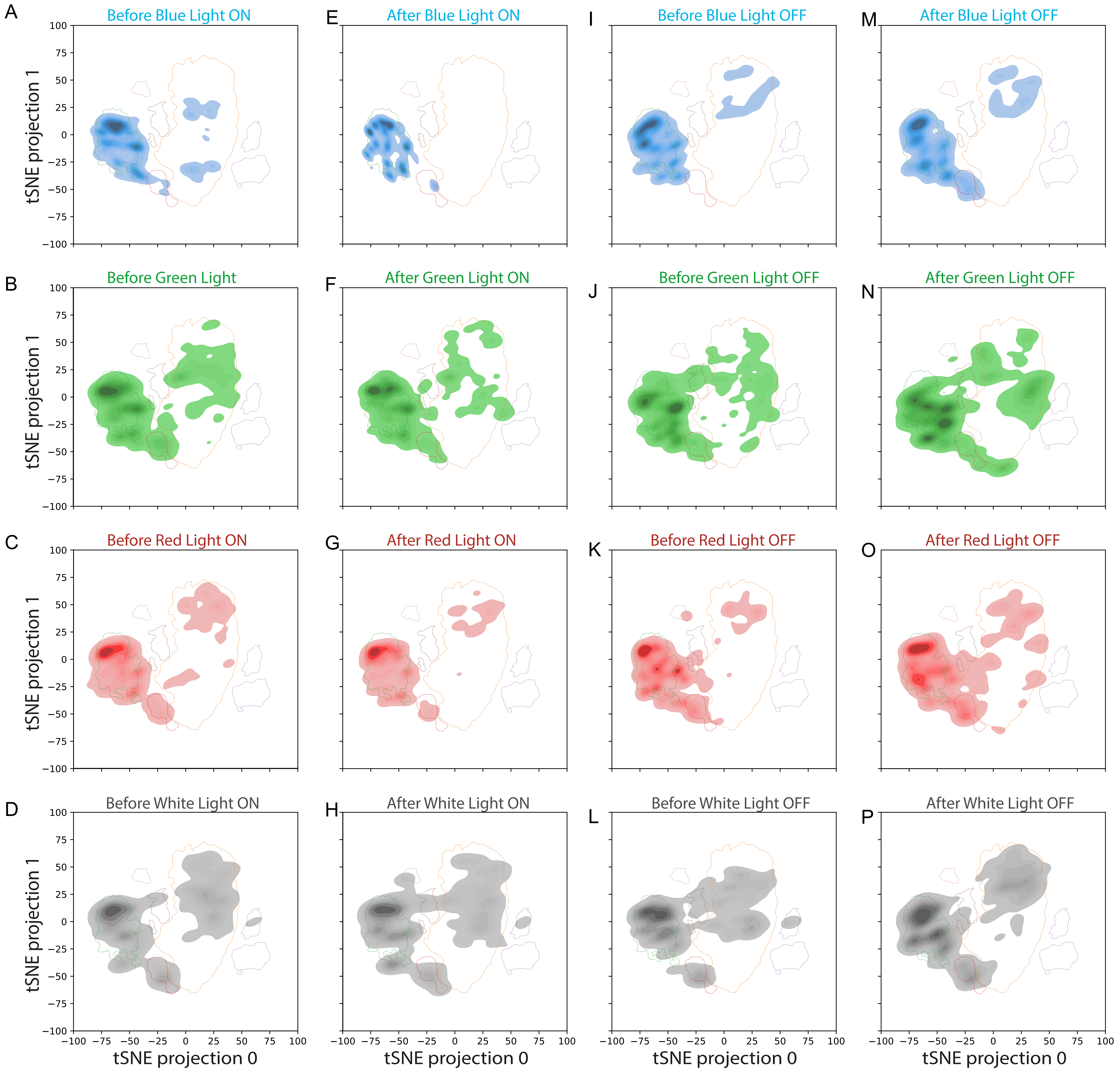
A
Before Blue Light ON
E
After Blue Light ON
I
Before Blue Light OFF
M
After Blue Light OFF
B
Before Green Light
F
After Green Light ON
J
Before Green Light OFF
N
After Green Light OFF
C
Before Red Light ON
G
After Red Light ON
K
Before Red Light OFF
O
After Red Light OFF
D
Before White Light ON
H
After White Light ON
L
Before White Light OFF
P
After White Light OFF
tSNE projection 1
tSNE projection 0
100
75
50
25
0
−25
−50
−75
−100

**Fig. S10.**

(A-P) This figure shows each color stimulus is shown individually, the data is the same as in Fig. 6. 2D density plots showing the change in the pattern of occupancy of the 2D behavioral space across different color light stimuli before and after the ON and OFF events. This figure shows each color stimulus individually, the data is the same as in Fig. 6. Boundaries of the 6 DBSCAN clusters are also shown on the plots.

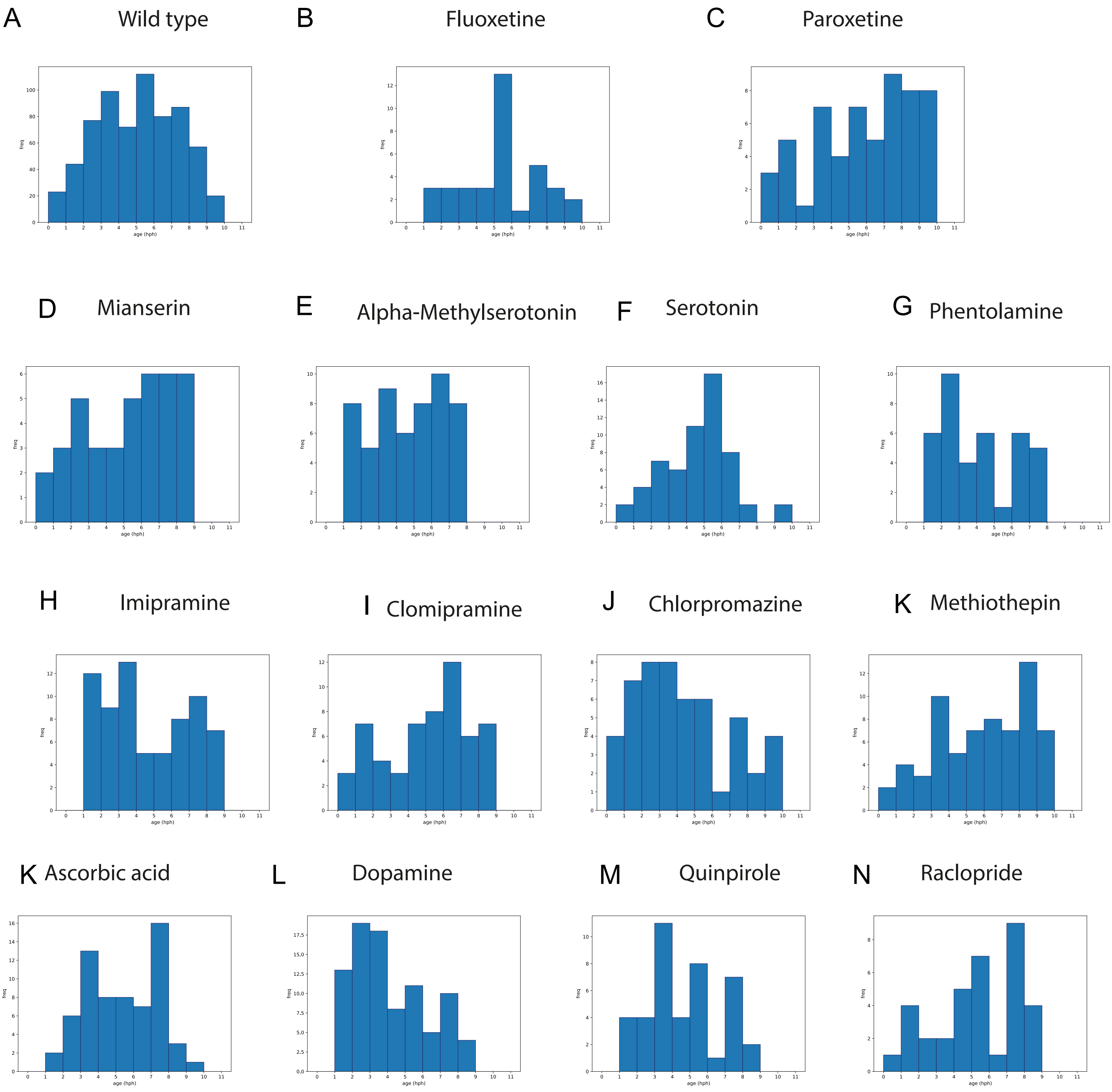
A Wild type
B Fluoxetine
C Paroxetine
D Mianserin
E Alpha-Methylserotonin
F Serotonin
G Phentolamine
H Imipramine
I Clomipramine
J Chlorpromazine
K Methiothepin
K Ascorbic acid
L Dopamine
M Quinpirole
N Raclopride
freq
age (hph)

**Fig. S11.**
(A-N) Histograms indicating the age distribution in hours post hatching (hph) for wild type and drug treated larvae.
